# Synaptic plasticity via receptor tyrosine kinase/G protein-coupled receptor crosstalk

**DOI:** 10.1101/2023.08.28.555210

**Authors:** Cristina Lao-Peregrin, Guoqing Xiang, Jihye Kim, Ipsit Srivastava, Alexandra B. Fall, Danielle M. Gerhard, Piia Kohtala, Daegeon Kim, Minseok Song, Mikel Garcia-Marcos, Joshua Levitz, Francis S. Lee

## Abstract

Cellular signaling involves a large repertoire of membrane receptors operating in overlapping spatiotemporal regimes and targeting many common intracellular effectors. However, both the molecular mechanisms and physiological roles of crosstalk between receptors, especially those from different superfamilies, are poorly understood. We find that the receptor tyrosine kinase (RTK), TrkB, and the G protein-coupled receptor (GPCR), metabotropic glutamate receptor 5 (mGluR5), together mediate a novel form of hippocampal synaptic plasticity in response to brain-derived neurotrophic factor (BDNF). Activated TrkB enhances constitutive mGluR5 activity to initiate a mode-switch that drives BDNF-dependent sustained, oscillatory Ca^2+^ signaling and enhanced MAP kinase activation. This crosstalk is mediated, in part, by synergy between Gβγ, released by TrkB, and Gα_q_-GTP, released by mGluR5, to enable a previously unidentified form of physiologically relevant RTK/GPCR crosstalk.

## Introduction

Cell signaling is based on a complex interplay between ensembles of receptors that sense extracellular signal dynamics and convert them into intracellular cascades that shape a multitude of functions ^1–3^. In the central nervous system, rapid synaptic transmission and its plasticity are mediated by ion channel-linked and G protein-coupled receptors (GPCRs) that sense neurotransmitters, such as glutamate ^4^. In contrast, growth factors like brain-derived neurotrophic factor (BDNF) typically signal via receptor tyrosine kinases (RTKs) on slower time scales to regulate the induction and expression of synaptic plasticity ^5^. In addition to their roles in normal brain function, both glutamate and BDNF signaling underlie many aspects of the pathophysiology and treatment of neurodevelopmental and neurodegenerative disorders ^6–8^. Despite both forms of signaling likely occurring coincidentally at excitatory synapses, little is known about how neurotrophin and neurotransmitter receptor classes work in concert to tune synaptic plasticity.

Tropomyosin-related kinase B (TrkB) and metabotropic glutamate receptor 5 (mGluR5) represent two of the most abundant, widely expressed membrane receptors in the brain with overlapping synaptic roles. TrkB is an RTK that senses BDNF, which drives autophosphorylation of intracellular tyrosine residues on the kinase domain to initiate intracellular signaling cascades through the binding of adaptor proteins ^9^. For example, the Shc adaptor protein links activated TrkB to two separate pathways: PI-3 kinase and MAP kinase signaling ^10^. In addition, phospholipase C-γ (PLC-γ) binds to phosphorylated TrkB and initiates IP_3_ production to drive release of intracellular Ca^2+^ stores. mGluR5 is a G_q_-coupled family C GPCR that uses heterotrimeric G proteins to initiate signaling cascades. Most prominently, mGluR5 activation leads to distinctive Ca^2+^ oscillations due to reversible protein kinase C-dependent phosphorylation of intracellular residues ^11–14^. In addition to its glutamate-evoked activity, under many circumstances, mGluR5 produces tonic glutamate-independent signaling ^15,16^.

In the hippocampus, BDNF-TrkB signaling contributes to tetanus-induced long term potentiation (LTP) ^17–19^, and exogenous BDNF application alone produces a chemical form of LTP (“BDNF-LTP”) ^20–23^. In contrast, hippocampal mGluR5 can contribute to electrical and chemical forms of either LTP or long-term depression (LTD), depending on the context ^4,24–29^. Both mGluR5 and TrkB signal via biochemical cascades and activate common downstream signaling effectors strongly implicated in synaptic plasticity, such as intracellular Ca^2+^ release and MAP kinase activation ^28,30^. These overlapping synaptic roles and signaling properties together raise the possibility that crosstalk between TrkB and mGluR5 may serve as a link between rapid neurotransmission and long-term plasticity.

It has previously been shown that GPCR activation can elicit signaling through transactivation of RTKs, such as the epidermal growth factor receptor (EGFR) or TrkB, through a variety of mechanisms including regulation of RTK ligand release or RTK phosphorylation following GPCR activation of the tyrosine kinase Src ^31–38^. However, minimal work has addressed the ability of a GPCR to alter the signaling response of an RTK to its native ligand ^39,40^. Using acute slice electrophysiology and cultured neuron imaging, we find that BDNF-TrkB-induced synaptic plasticity is dependent on constitutive, coincident dendritic mGluR5 signaling. Using a battery of assays in cultured cells, we reveal that mGluR5 co-expression or positive allosteric modulation dramatically boosts the BDNF sensitivity and response duration of TrkB in terms of both intracellular Ca^2+^ release and MAP kinase pathway activation. This crosstalk is mediated by constitutive G protein activation by mGluR5 and non-canonical activation of G protein signaling by TrkB which, together, enables cooperative activation of downstream effectors such as phospholipase C-β (PLC-β). This G protein-dependent crosstalk mechanism underlies the mGluR5-dependence of BDNF-induced synaptic plasticity and occurs for a range of RTKs and GPCRs, indicating that this is a general mode of RTK/GPCR synergy that is relevant across physiological systems.

## Results

### mGluR5 activity is required for BDNF-induced synaptic plasticity

Given that both TrkB and mGluR5 can drive certain forms of LTP in the hippocampal CA3-CA1 synapse, we asked if these receptors functionally interact in this context. We performed field recordings from adult mouse hippocampal slices with exogenous BDNF application, as done previously ^20–23^. Application of BDNF for 30 min produced a dose-dependent increase in fEPSP slope that persisted for an hour after washout (**Fig. S1A**). This effect was blocked by the selective TrkB antagonist ANA-12 ^41^ (**Fig. S1B**). Throughout this study, we refer to this chemical form of long-term synaptic plasticity as “BDNF-LTP”. As a first test for a potential contribution of mGluR5 to this form of plasticity, we asked if pharmacological inhibition of mGluR5 would alter BDNF-LTP induction or expression. Strikingly, when slices were pre-treated and maintained in the mGluR5 negative allosteric modulator MPEP (40 μM) during BDNF (100 ng/mL) application, induction of BDNF-LTP was largely diminished (**Fig. 1A, B**). Application of MPEP in the absence of BDNF had no effect on fEPSP slope (**Fig. S1C**). MPEP treatment following BDNF-induced potentiation also had no effect (**Fig. S1D**), indicating that mGluR5 activation is required for BDNF-LTP induction, but not for expression or maintenance. Furthermore, BDNF-LTP was not associated with changes in paired pulse ratio (**Fig. S1E**), as shown previously ^22^, in line with a post-synaptic mechanism. Consistent with pharmacological blockade of mGluR5, post-development conditional knockout (KO) of mGluR5 in CA1 neurons via virally delivered Cre-recombinase (**Fig. S1F**) prevented BDNF-LTP (**Fig. 1C, D**). Importantly, CA1 knockout of mGluR5 also strongly diminished sensitivity to the group I mGluR agonist, dihydroxyphenylglycine (DHPG) (**Fig. S1G, H**), but did not alter basal synaptic strength as assessed with a titration of stimulus intensity (**Fig. S1I**).

**Figure 1.**
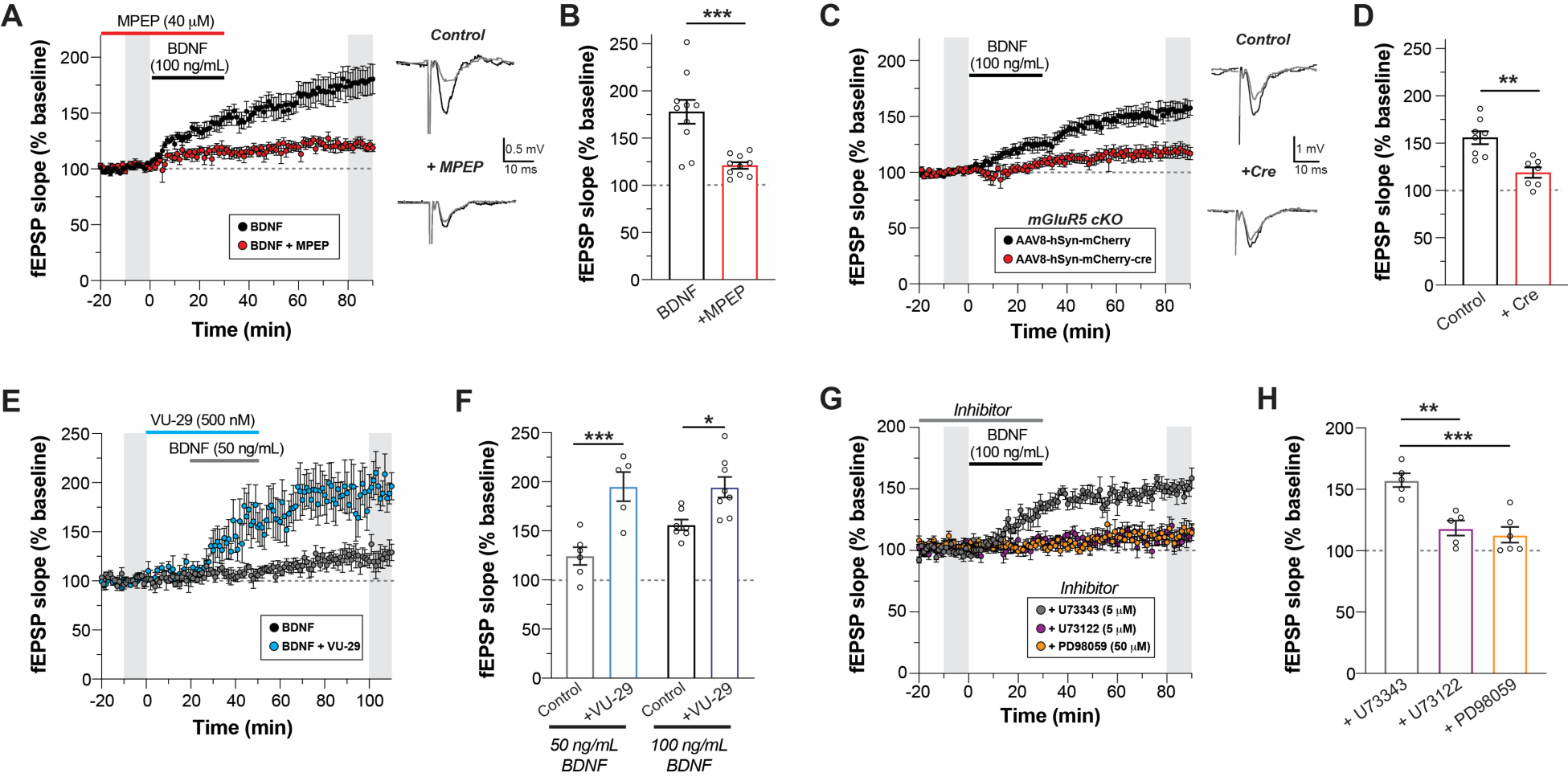
BDNF-induced LTP is dependent on mGluR5 and enhanced by a mGluR5 allosteric modulator. **(A)** fEPSP slope time course showing that blockade of mGluR5 with MPEP prevents BDNF-induced LTP. Grey bars show regions averaged for baseline and post-BDNF values in (B). Right, representative fEPSP traces recorded during basal and after 60 min of BDNF perfusion. **(B)** Summary bar graph showing a lack of BDNF-induced potentiation in the presence of MPEP. **(C-D)** Conditional KO of mGluR5 in CA1 pyramidal neurons impairs BDNF-LTP compared to control slices. **(E-F)** Co-application of the mGluR5 PAM VU-29 enhances LTP induced by low dose (50 ng/mL) or high dose (100 ng/mL) BDNF. **(G-H)** BDNF-induced LTP is blocked by an ERK inhibitor (PD98059, 50 μM) or a PLC inhibitor (U-73122, 5 μM), but not an inactive PLC inhibitor analog (U-73343, 5 μM). For (B), (D), (F), (H): individual points represent independent slices taken from distinct mice. For (B) and (D), unpaired t-test is used. For (F) and (H), one-way ANOVA with Sidak’s multiple comparisons is used. All data shown as mean ± SEM; * P < 0.05, ** P < 0.01, *** P < 0.001. See also Figures S1 and S2.

Given that mGluR5 activity is required for BDNF-LTP induction, we hypothesized that positive allosteric modulators (PAMs) of mGluR5 may enhance BDNF-induced potentiation. A prior study showed that the mGluR5 PAM VU-29 can potentiate tetanus-induced LTP ^26^, which also involves BDNF-TrkB signaling ^30^. Consistent with this study, VU-29 did not alter baseline synaptic transmission (**Fig. S1J**). However, co-application of VU-29 with a lower dose of BDNF (50 ng/mL) dramatically enhanced LTP (**Fig. 1E, F**). VU-29 was also able to modestly enhance the LTP induced by a higher BDNF dose (100 ng/mL) (**Fig. 1F**). The effects of VU-29 further indicate that synergistic interplay between TrkB and mGluR5 can drive synaptic strengthening. For further insight into the signaling mechanisms underlying mGluR5-dependent BDNF-LTP, we assessed the contributions of extracellularly regulated kinase (ERK) and phospholipase C (PLC), two key signaling molecules previously implicated in BDNF-mediated plasticity ^42,43^. Application of either PD98059, an ERK1/2 inhibitor, or U73122, a PLC inhibitor, had no effect on basal fEPSP slope (**Fig. S1K, L**). However, co-application of PD98059 or U73122 with BDNF prevented BDNF-LTP (**Fig. 1G, H**), suggesting that both ERK and PLC signaling are required for the induction of BDNF-LTP.

In addition to its ability to tune the electrical properties of glutamatergic synapses, BDNF can also produce morphological changes in dendritic spines ^44–46^. This raises the possibility that mGluR5 also contributes to structural forms of BDNF-induced synaptic plasticity. To assess whether TrkB and mGluR5 are co-localized in such a way that they may co-regulate spine growth, we visualized both endogenous receptors in primary cultured hippocampal neurons using immunocytochemistry and scanning confocal microscopy (**Fig. 2A**). Both TrkB and mGluR5 puncta were observed along the dendritic shaft and spines, including a substantial co-localized population within the same spine or shaft region (**Fig. 2A**). In addition, mGluR5 was detected in TrkB immunoprecipitates from hippocampal neurons, as assessed via western blot (**Fig. S2A**). These results provide evidence that mGluR5 and TrkB compartmentalize together within signaling microdomains, providing a basis for functional crosstalk via access to an overlapping pool of signaling targets.

**Figure 2.**
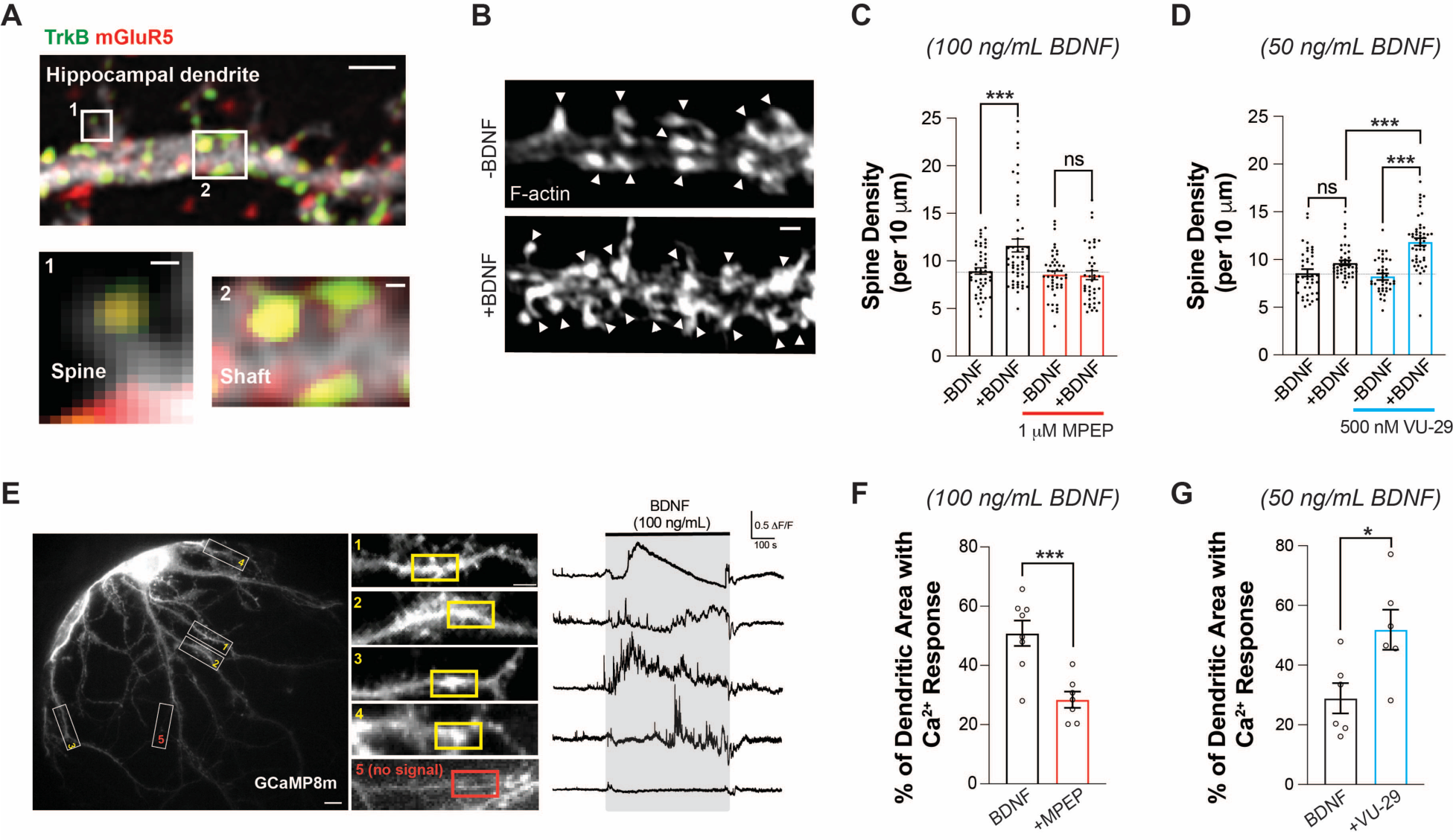
BDNF-induced dendritic spine growth and calcium signaling are dependent on mGluR5. **(A)** Confocal image of fixed hippocampal neuron showing anti-TrkB and anti-mGluR5, including colocalization in dendrites (Scale bars 1 µm dendrite; 0.1 µm spine, shaft). 49.3 ± 6.6 % of spines showed co-expression of TrkB and mGluR5; Pearson’s correlation coefficient between TrkB and mGluR5 in dendritic shaft (top 10% of pixels) = 0.49 ± 0.05 (n = 12 neurons). **(B)** Representative images showing BDNF-induced (100 ng/mL) spine density increases in DIV 21 hippocampal neurons. Arrowheads denote spine heads enriched with F-actin. **(C-D)** Bar graphs summarizing the BDNF-induced increases in spine density. MPEP blocks 100 ng/mL BDNF-induced spine density increase (C) while VU-29 potentiates the increase in spine density induced by low dose 50 ng/mL BDNF (D). **(E)** Representative image of GCaMP8m-expressing hippocampal neuron (left, scale bar 10 µm), with snapshots of dendrites (white rectangle) taken in the presence of 100 ng/mL BDNF (middle, scale bar 5 µm). Right, representative trace from 10 µm ROIs with Ca^2+^ response (yellow rectangle) or without Ca^2+^ response (blue rectangle) from each dendrite. **(F)** Bar graph showing that 1 μM MPEP co-application significantly decreases the percentage of total dendritic area with BDNF-induced Ca^2+^ responses. (**G**) Bar graph showing that co-application of low dose (50 ng/mL) BDNF with 500 nM VU-29 leads to an increased percentage of total dendritic area with Ca^2+^ responses. For (C), (D), (F), (G): individual points represent separate neurons, taken from at least 3 separate culture preparations. For (C) and (D), one-way ANOVA with Tukey’s multiple comparisons is used. For (F) and (G), unpaired t-test is used. All data shown as mean ± SEM, * P < 0.05, *** P < 0.001. See also Figure S2.

To assess BDNF-induced spine growth, we acutely added BDNF for 30 min in mature primary hippocampal neurons. We observed a dose-dependent 20-30% increase in spine density (**Fig. 2B, C; Fig. S2B**), which was prevented by pre-treatment with ANA-12 (**Fig. S2C**). As was seen with field recordings in acute slices, BDNF-induced spine density increase was not observed in the presence of MPEP (**Fig. 2C**), indicating that BDNF-induced structural plasticity is also dependent on mGluR5. Furthermore, VU-29 enhanced BDNF-induced structural plasticity (**Fig. 2D**) while PLC or ERK inhibition prevented BDNF-induced spine growth (**Fig. S2D, E**). Together, these data show that electrical and structural forms of BDNF-driven plasticity share common mGluR5-dependent properties.

### mGluR5 co-expression and allosteric modulation controls BDNF-TrkB signaling dynamics

Based on the PLC-dependence of BDNF-LTP in our study and prior studies implicating intracellular Ca^2+^ stores in BDNF-LTP ^47^, we assessed the dendritic Ca^2+^ responses to BDNF application using the genetically-encoded fluorescent sensor GCaMP8m in primary hippocampal neuronal cultures. In the presence of tetrodotoxin (TTX) to prevent action potential firing and associated neurotransmitter release, sporadic Ca^2+^ events were observed in dendrites. Application of BDNF led to a substantial increase in the amount of dendritic Ca^2+^ events, which showed a wide variety of temporal dynamics and were confined within ∼5-30 µm dendritic patches (**Fig. 2E**). Pre-application of MPEP substantially decreased the proportion of dendritic area showing BDNF responses (**Fig. 2F; Fig. S2F**), in line with its inhibitory effect on BDNF-LTP and spine growth. Furthermore, application of VU-29 enhanced the response to a lower dose BDNF (50 ng/mL) (**Fig. 2G; Fig. S2G**). These results show that mGluR5 activity contributes to the initial dendritic response to BDNF, motivating a subsequent focus on how mGluR5 shapes acute BDNF-TrkB Ca^2+^ signaling dynamics.

To further probe the signaling that underlies the mGluR5-dependence of BDNF-TrkB synaptic plasticity, we turned to a simplified HEK 293 cell context. In a stable cell line expressing TrkB (“HEK 293-TrkB”) ^48^, BDNF produced dose-dependent Ca^2+^ responses in the form of a single transient of 50-200 sec duration, consistent with the established ability of TrkB to activate PLC-γ ^49^ (**Fig. 3A; Fig. S3A-E**). In contrast to RTKs, Gα_q_-coupled GPCRs, such as mGluR5, signal via the PLC-β family ^50^. HEK 293 cells transiently transfected with mGluR5 responded to glutamate application with oscillatory Ca^2+^ responses with event durations of ∼30 sec and were insensitive to BDNF (**Fig. 3B; Fig. S3F, G**). mGluR5-induced Ca^2+^ oscillations are due to reversible protein kinase C (PKC)-mediated phosphorylation of receptor C-terminal domain residues ^11,13,14^. Based on initial event duration (**Fig. S3G**), responses were classified as “slow Ca^2+^ wave” or “Ca^2+^ oscillations” with the former associated with TrkB and the latter with mGluR5.

**Figure 3.**
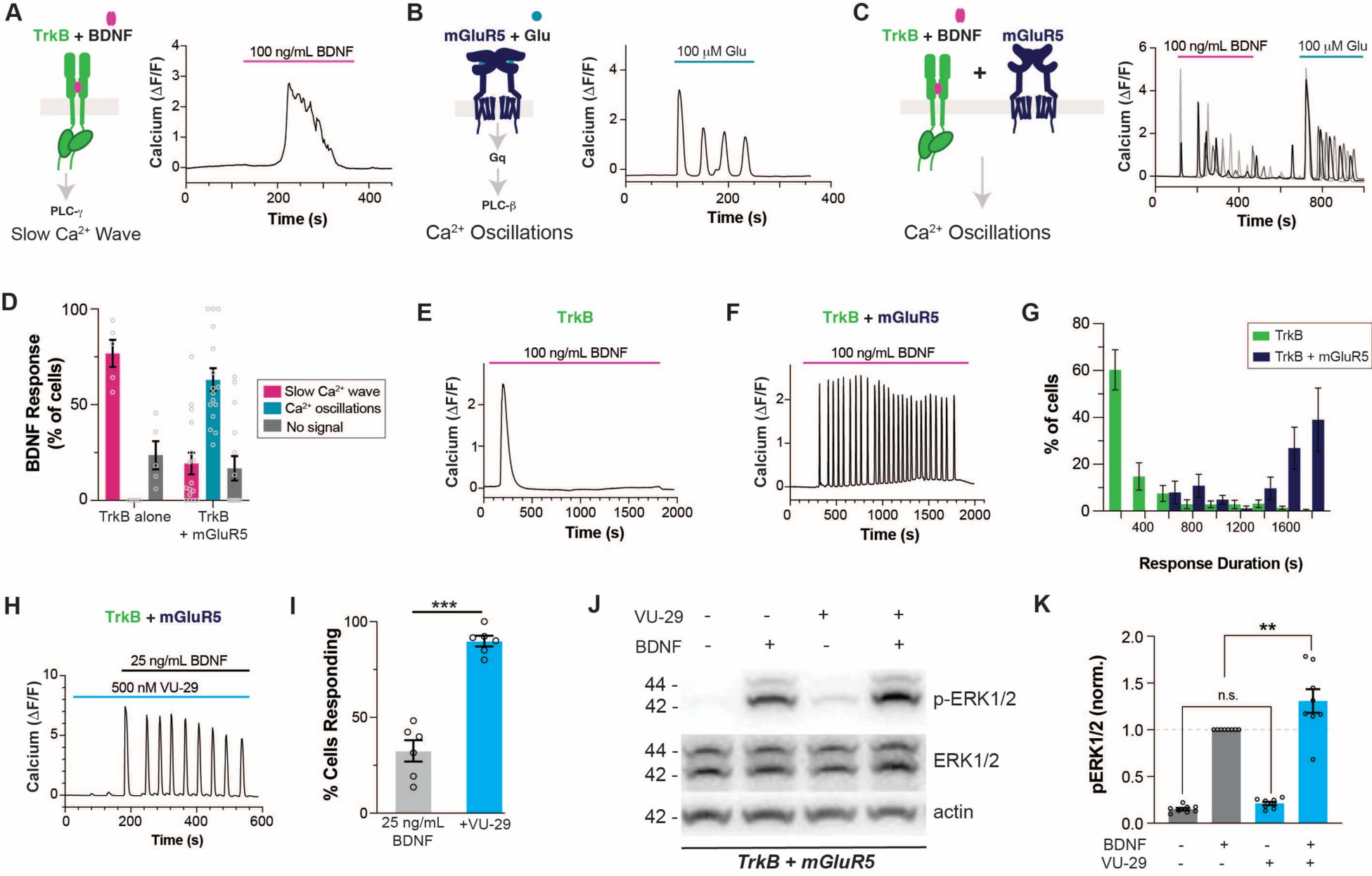
BDNF-mediated TrkB activation produces mGluR5-dependent Ca^2+^ oscillations in HEK 293 cells. **(A-C)** Representative traces showing intracellular Ca^2+^ responses to TrkB activation by BDNF (A), mGluR5 activation by glutamate (B) and TrkB activation by BDNF in mGluR5 co-expressing cells (C). **(D)** Distribution of BDNF responses in the absence or presence of mGluR5 co-expression. Only cells showing a response to glutamate are analyzed in the TrkB + mGluR5 condition. **(E-G)** Representative traces showing Ca^2+^ responses to extended 30 min BDNF application in cells expressing TrkB (E) or TrkB and mGluR5 (F), with summary histogram (G) showing the distribution of response durations. **(H-I)** Co-application of the mGluR5 PAM VU-29 enhances the response to low dose BDNF, as seen in a representative cell (H) and a summary bar graph of the percentage of cells responding to BDNF (I). Only cells responding to glutamate were including in the bar graph in (I). **(J-K)** Representative western blot (J) and quantification (K) of BDNF-induced ERK activation (p-ERK/ERK ratio at 15 min) in HEK 293-TrkB cells co-expressing mGluR5, showing an enhanced response in the presence of 500 nM VU-29. For (D) and (I), points represent values from individual movies taken from distinct coverslips. For (K), individual points represent value from individual blots. For all conditions, data comes from at least 3 separate cell preparations. Unpaired t-test for (I); One-way ANOVA with Tukey’s multiple comparisons for (K). All data shown as mean ± SEM; ** P < 0.01, *** P < 0.001. See also Figure S3 and S4.

Strikingly, upon co-expression of mGluR5 in HEK 293-TrkB cells, BDNF elicited Ca^2+^ oscillations in ∼60% of glutamate-sensitive cells while only ∼20% produced a slow Ca^2+^ wave response (**Fig. 3C, D**). This suggests a mode-switch in BDNF-induced Ca^2+^ signaling upon mGluR5 co-expression. In addition to BDNF, neurotrophin-3 (NT-3) and neurotrophin-4 (NT-4), which also activate TrkB, produced mGluR5-dependent Ca^2+^ oscillations (**Fig. S4A-D**). BDNF-induced Ca^2+^ oscillations were observed with application of low dose (25 ng/mL) (**Fig. S4E, F**) and high dose (100 ng/mL) of BDNF (**Fig. 3C, D**). Notably, mGluR5 co-expression slightly decreased the amplitude (**Fig. S4G**) and substantially decreased the latency of BDNF responses (**Fig. S4H**), demonstrating a robust acceleration of TrkB signaling. When BDNF was applied continuously for 30 min, as was done for synaptic plasticity induction (**Fig. 1, 2**), TrkB alone produced a single slow Ca^2+^ wave (**Fig. 3E**) while mGluR5 co-expression enabled Ca^2+^ oscillations that persisted for the entire BDNF application time (**Fig. 3F, G**). Similarly, mGluR5 co-expression increased the amplitude and duration of ERK phosphorylation following BDNF treatment in HEK 293-TrkB cells (**Fig. S4I, J**). We also found that application of the mGluR5 PAM VU-29 did not directly activate mGluR5 Ca^2+^ signaling (**Fig. S4K, L**) but dramatically increased the proportion of cells responding to low dose BDNF (25 ng/mL) with Ca^2+^ oscillations (**Fig. 3H, I; Fig. S4M**), showing that mGluR5 can effectively sensitize TrkB to its endogenous ligand. With a high dose of BDNF, VU-29 enhanced the proportion of cells responding to >90% (**Fig. S4N**). VU-29 also increased the amplitude (**Fig. S4G**), decreased the latency (**Fig. S4H**), and increased the Ca^2+^ oscillation frequency of BDNF responses (**Fig. S4O**). Lastly, co-application of VU-29 increased the extent of ERK phosphorylation following low dose (25 ng/mL) BDNF application in HEK 293-TrkB cells co-expressing mGluR5 (**Fig. 3J, K**). Together, these data indicate that mGluR5 co-expression and positive allosteric modulation can amplify the TrkB response to BDNF, which may underlie the mGluR5-dependence of BDNF-LTP induction.

### Molecular mechanisms of TrkB/mGluR5 signaling crosstalk

We next investigated the molecular mechanisms that mediate TrkB/mGluR5 signaling crosstalk. Like many GPCRs, mGluR5 has previously been shown to display constitutive activity ^15^, which may explain its contribution to the BDNF response in the absence of glutamate application. However, the relatively high affinity of mGluR5 for glutamate makes it hard to rule out a role for ambient glutamate or for BDNF-induced glutamate release as a driver of Ca^2+^ oscillations. To clarify this, we removed the extracellular domains of mGluR5, including the glutamate binding domain, to produce “mGluR5-ι1ECD” ^51^. Co-expressing mGluR5-ι1ECD with TrkB did not prevent BDNF-induced Ca^2+^ oscillations (**Fig. 4A, B**). Importantly, prior work has shown that mGluR5-ι1ECD still can couple to G proteins and shows constitutive activity ^51^. Indeed, VU0360172, an mGluR5 allosteric agonist, still elicited robust Ca^2+^ oscillations with this construct (**Fig. 4A**). We then probed if mGluR5 activation is required for BDNF-induced Ca^2+^ oscillations by using MPEP, which fully abolished BDNF-induced Ca^2+^ oscillations, indicating that mGluR5 activation is required (**Fig. 4B; Fig. S5A**). Importantly, MPEP did not alter BDNF Ca^2+^ responses in cells expressing TrkB but not mGluR5 (**Fig. S5B**). MPEP treatment also blocked slow Ca^2+^ wave responses (**Fig. S5A**) and decreased the ERK phosphorylation response to BDNF in cells co-expressing TrkB and mGluR5 (**Fig. S5C, D**). Furthermore, introduction of a mutation to a conserved intracellular loop 3 residue ^52^ (F768D) to prevent mGluR5 from coupling with G proteins, also prevented BDNF-induced Ca^2+^ oscillations (**Fig. 4B; Fig. S5E, F**) without altering mGluR5 surface levels (**Fig. S5G**). Together, these experiments show that TrkB/mGluR5 signaling crosstalk is dependent on mGluR5/G protein coupling but is independent of glutamate binding to mGluR5.

**Figure 4.**
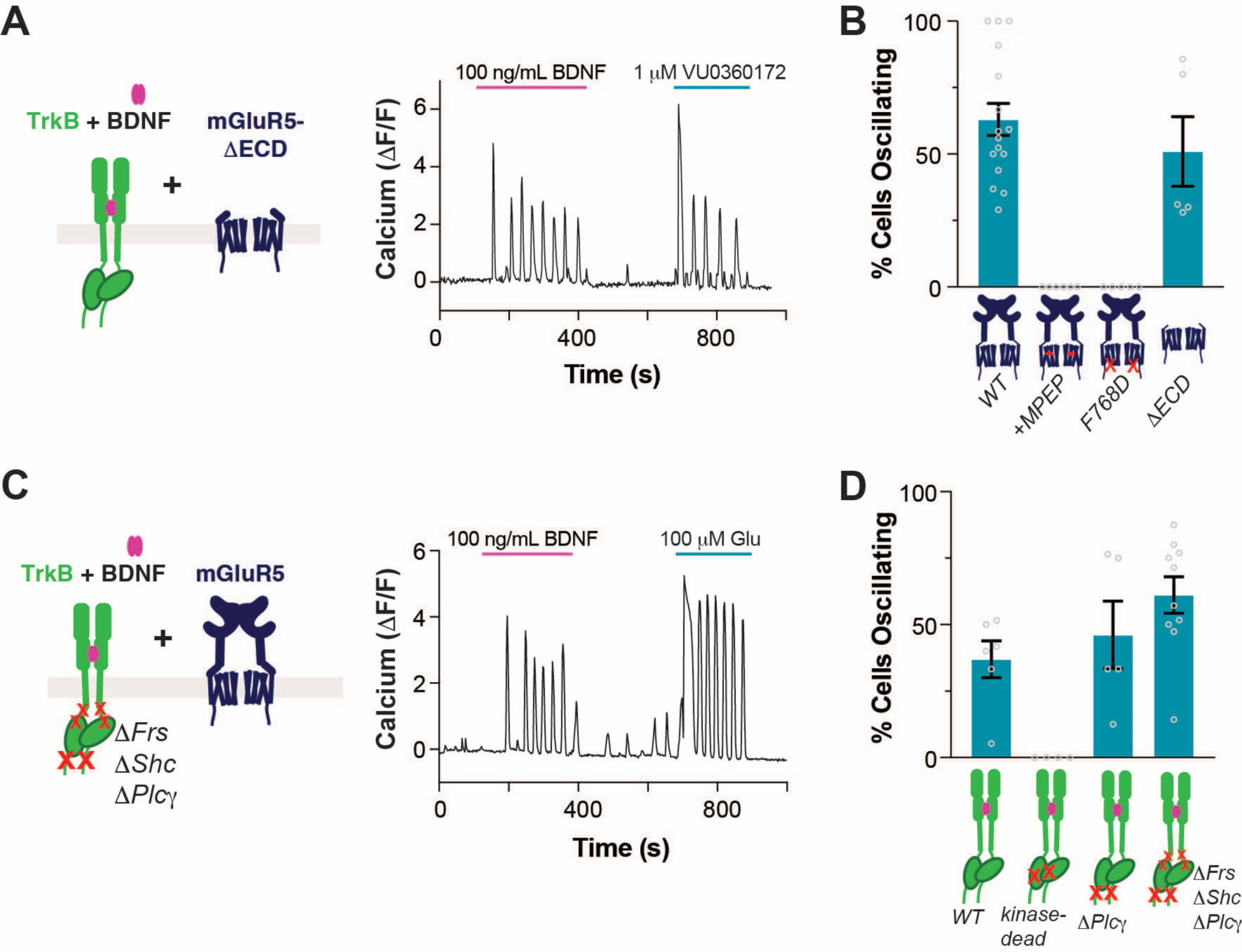
Molecular determinants of BDNF-induced Ca^2+^ oscillations. **(A)** Representative trace showing BDNF-induced Ca^2+^ oscillations in cells co-expressing TrkB and mGluR5-ΔECD. Ca^2+^ oscillations are also produced with VU0360172, a mGluR5 allosteric agonist. **(B)** Summary bar graph showing lack of BDNF-induced Ca^2+^ oscillations when mGluR5 is blocked by MPEP or when the F768D mutation is introduced, but not when the ECD is removed. Only cells responding to glutamate or VU0360172 (for mGluR5-ΔECD) were analyzed. **(C)** Representative trace showing BDNF-induced Ca^2+^ oscillations in cells co-expressing mGluR5 and TrkB-ΔFrs-ΔShc-ΔPLCγ. **(D)** Summary bar graph showing lack of BDNF-induced Ca^2+^ oscillations with kinase-dead TrkB (K571N), but clear oscillations for TrkB-ΔPLCγ and TrkB-ΔFrs-ΔShc-ΔPLCγ. Only cells showing a response to glutamate were analyzed. Points in (B) and (D) represent values from individual movies taken from distinct coverslips. For all conditions, data comes from at least 3 separate cell preparations. All data shown as mean ± SEM. See also Figure S5.

To probe the determinants of TrkB/mGluR5 crosstalk at the level of TrkB via perturbations directly to TrkB, we turned to transient co-transfection of TrkB and mGluR5, which also enabled BDNF-induced, mGluR5-dependent Ca^2+^ oscillations (**Fig. 4D**). To determine if TrkB kinase activity is required for Ca^2+^ oscillations, we tested the TrkB-K571N “kinase-dead” mutant ^53^. Introduction of this mutation prevented BDNF-induced Ca^2+^ responses in the presence or absence of mGluR5 (**Fig. 4D; Fig. S5H-J**). In contrast, removal of the PLC-γ recruitment site alone (“TrkB-ΔPLCγ”) or in combination with the Shc and FRS2 sites (“TrkB-ΔFrs-ΔShc-ΔPLCγ”) did not prevent BDNF-induced Ca^2+^ oscillations in cells co-expressing mGluR5 (**Fig. 4C, D; Fig. S5I**), despite preventing Ca^2+^ responses in cells only expressing TrkB (**Fig. S5J**). As a key control, we showed that both the TrkB kinase dead and the TrkB-ΔFrs-ΔShc-ΔPLCγ are unable to activate the ERK pathway, while the TrkB-ΔPLCγ maintained a weak BDNF-induced ERK response (**Fig. S5K, L**). Furthermore, all TrkB mutants showed clear surface expression, with TrkB-ΔFrs-ΔShc-ΔPLCγ showing increased surface expression compared to wild-type TrkB (**Fig. S5M**). These results indicate that BDNF-induced Ca^2+^ oscillations are not dependent on canonical TrkB signaling mechanisms via Shc, FRS2, or PLC-γ pathways, but they do require tyrosine kinase activity.

The ability of BDNF to produce mGluR5-dependent Ca^2+^ oscillations via the TrkB-ΔFrs-ΔShc-ΔPLCγ construct, which lacks canonical mechanisms for downstream signaling, motivated us to investigate alternative pathways. A variety of RTKs have been shown to couple to heterotrimeric G proteins via non-canonical mechanisms that nonetheless rely on kinase activity ^54–58^. Furthermore, a recent study showed that TrkB is highly sensitive to knockout of Gα_i1_ and Gα_i3_ G proteins, which dramatically reduced BDNF-induced signaling in hippocampal neurons ^59^. We hypothesized that upon BDNF binding, TrkB can promote G_i/o_ signaling, which then synergizes with G_q/11_ proteins tonically activated by mGluR5 (**Fig. 5A**). Informed by a model proposed to explain the crosstalk between Gα_i/o_-coupled GPCRs and G_q/11_-coupled GPCRs ^60,61^, we reasoned that Gβγ subunits released by Gα_i/o_ under the control of TrkB and Gα_q_-GTP promoted by mGluR5 would synergize by simultaneously binding to PLC-β to produce Ca^2+^ oscillations. Consistent with this idea and with prior work showing that group I mGluR signaling can be promoted by Gα_i/o_-coupled receptors ^62,63^, we found that activation of Gα_i/o_ signaling upon stimulation of different GPCRs (mu-opioid receptor, MOR, or gamma-aminobutyric acid B receptor, GABA_β_R) led to mGluR5-dependent Ca^2+^ oscillations in the absence of glutamate (**Fig. S6A, B**). The similarities of these responses with those elicited by BDNF (**Fig. 3**) motivated us to test the effect of G protein perturbations on the interplay between TrkB and mGluR5.

**Figure 5.**
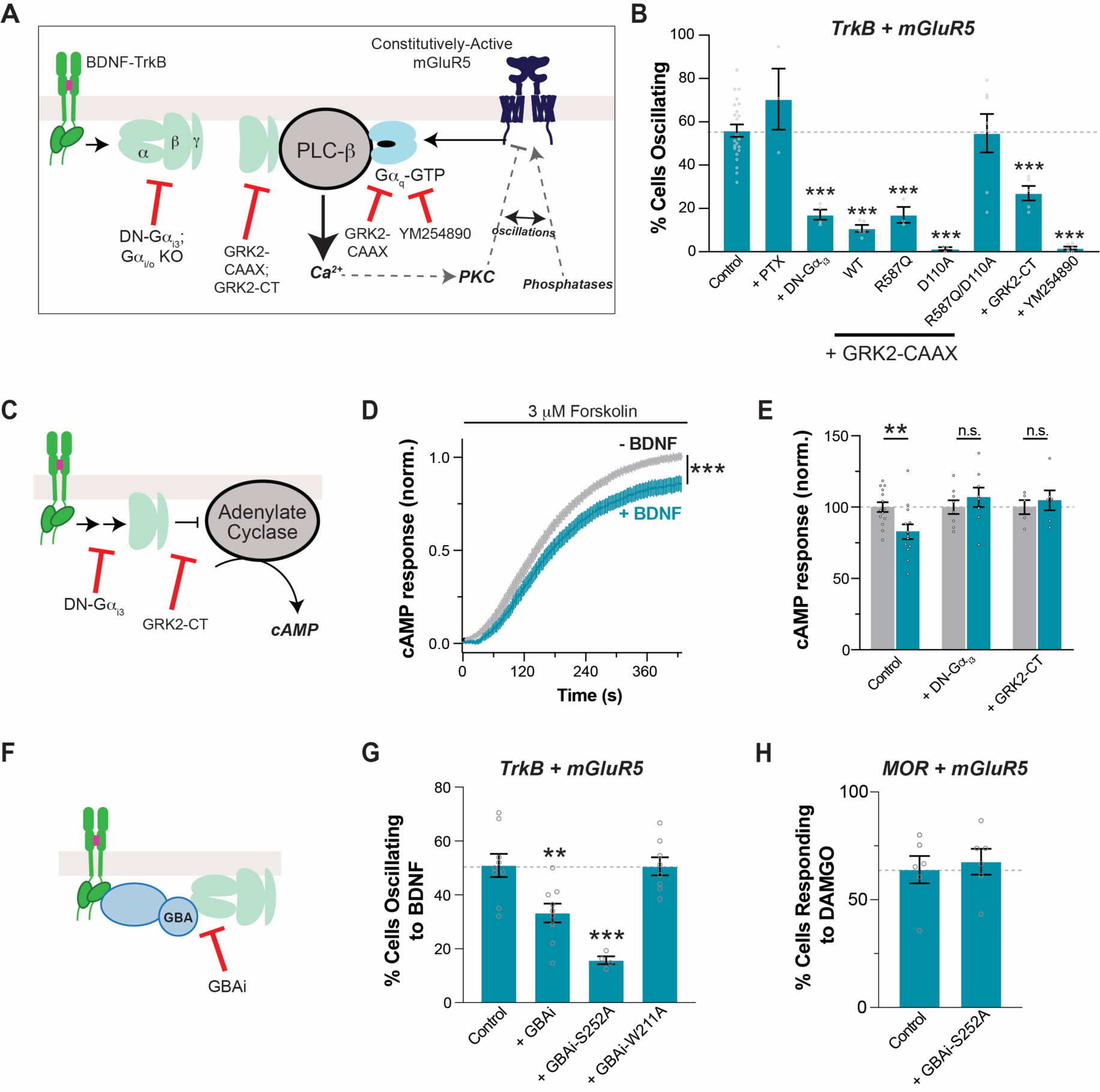
G protein-dependence of TrkB/mGluR5 crosstalk. **(A)** Schematic of the proposed “G protein synergy” mechanism underlying TrkB/mGluR5 crosstalk. **(B)** Summary bar graph showing effects of G protein perturbations on the efficiency of BDNF-induced Ca^2+^ oscillations in cells expressing TrkB and mGluR5. “PTX” = pertussis toxin; “DN-Gα_i3_” = Gα_i3_-G203A; “GRK2-CAAX” = membrane-tethered GRK2; R587Q impairs Gβγ binding; D110A impairs Gα_q_ binding; “GRK2-CT” = isolated, membrane-tethered PH domain of GRK2; YM-254890 (20 μM) = Gα_q_ blocker. **(C-E)** Schematic of cAMP signaling (C) with average traces (D) and summary bar graph (E) showing that TrkB activation by BDNF inhibits cAMP production in a G protein dependent manner. **(F-H)** Schematic of GBAi inhibition of G protein coupling (F) with summary bar graphs showing that GBAi over-expression decreases TrkB/mGluR5 crosstalk (G) but not MOR/mGluR5 crosstalk (H). For (B) and (F-H), only cells showing a response to glutamate were analyzed except for the YM-254890 condition, in which only cells responding to BDNF were included. Points represent values from individual movies taken from distinct cover slips (≥3 separate cell preparations per condition). All conditions were compared to the control group using one-way ANOVA with Dunnett’s multiple comparisons. All data shown as mean ± SEM; ** P<0.01; *** P < 0.001. See also Figure S6.

First, we found that co-expression of a dominant negative Gα_i3_ mutant, G203A (“DN-Gα_i3_”), that prevents the dissociation of Gβγ from Gα ^64–66^ dramatically reduced the proportion of cells showing mGluR5-dependent Ca^2+^ oscillations in response to BDNF in HEK 293-TrkB cells (**Fig. 5B**). Importantly, DN-Gα_i3_ did not alter BDNF responses in cells expressing TrkB alone or mGluR5-mediated glutamate responses (**Fig. S6C, D**). This dependence on G_i/o_ signaling was also corroborated by the observation of a large reduction in DAMGO-induced Ca^2+^ oscillations in mGluR5 and MOR-expressing cells upon expression of DN-Gα_i3_ (**Fig. S6E**). However, BDNF and DAMGO were affected differently by pertussis toxin (PTX), which modifies the C-terminus of Gα subunits in the G_i/o_ family ^67^. While PTX completely blunted DAMGO-induced mGluR5-dependent Ca^2+^ oscillations (**Fig. S6E**), as expected for a GPCR-mediated mechanism, it had no effect on BDNF-induced oscillations (**Fig. 5B**). The latter lends confidence to the idea that BDNF exerts its action through a non-canonical, RTK-dependent G_i/o_-Gβγ signaling that is PTX-insensitive ^57^ rather than through the activation of a GPCR intermediate.

We then co-expressed a membrane-tethered version of GPCR kinase 2 (“GRK2-CAAX”), which can bind and sequester both Gβγ and Gα_q_-GTP ^68,69^. Indeed, GRK2-CAAX co-expression reduced the proportion of cells responding to BDNF with Ca^2+^ oscillations (**Fig. 5B**), without altering BDNF responses in cells expressing TrkB alone (**Fig. S6C**) or mGluR5 glutamate responses (**Fig. S6D**). Mutation to both the Gβγ (R587Q) and Gα_q_ (D110A) binding sites on GRK2 were required to rescue efficient TrkB/mGluR5 crosstalk (**Fig. 5B**). Notably, the isolated GRK2 C-terminus (“GRK2-CT”), which contains only the Gβγ binding site, was also able to blunt both TrkB/mGluR5 (**Fig. 5B**) and MOR/mGluR5 crosstalk (**Fig. S6E**) without altering BDNF responses in cells expressing TrkB alone (**Fig. S6C**) or mGluR5 glutamate responses (**Fig. S6D**). The Gα_q_ inhibitor YM-254890 was able to block TrkB/mGluR5 crosstalk (**Fig. 5B**) and mGluR5 glutamate responses (**Fig. S6D**), but not slow wave Ca^2+^ responses from TrkB (**Fig. S6C**). Together, these data are consistent with a working “G protein synergy” model that is sensitive to blockade of either TrkB-induced Gβγ release or mGluR5-induced Gα_q_ activation (**Fig. 5A**).

Based on these results, we further probed the potential Gα_i/o_-coupling of TrkB. First, we tested if BDNF-mediated activation of TrkB can lead to similar downstream effects as Gα_i/o_-coupled GPCRs. We turned to cAMP imaging using the “cAMPr” sensor ^70^. Upon forskolin application to stimulate adenylate cyclase activity, Gα_i/o_-coupled receptor agonism typically decreases the enhancement of cAMP levels, as seen with MOR activation by DAMGO (**Fig. S6F, G**). BDNF application decreased the forskolin response in TrkB-expressing cells (**Fig. 5C-E**). Consistent with a G protein-dependent mechanism, BDNF-induced cAMP inhibition was not seen upon co-expression of DN-Gα_i3_ or GRK2-CT (**Fig. 5E**). However, it is worth noting that the inhibition of cAMP levels by TrkB (∼15%) was substantially smaller than that by MOR (∼50%), indicating that TrkB is a much less efficient activator of G proteins. Consistent with this apparent functional interaction between TrkB and Gα_i/o_ proteins, TrkB was able to co-IP Gα_i3_ both in HEK 293 cells and neurons (**Fig. S6H**), as has previously been shown ^59^.

We then asked if the family of G protein regulators that contain a G protein-Binding-and-Activating (“GBA”) motif mediate the effects of TrkB on the crosstalk with mGluR5. This family of broadly expressed guanine nucleotide exchange factors (GEFs), including proteins like Girdin/GIV and DAPLE, has been shown to drive nucleotide exchange and Gβγ release from G protein heterotrimers ^57,71–76^, including in response to RTK activation ^56,57,72–74,76,77^. To determine the necessity of direct involvement of G proteins by TrkB to crosstalk with mGluR5, we turned to a previously reported tool, termed GBAi ^78,79^. This is an engineered synthetic protein based on Gα that binds with high affinity to GBA motifs, but not to other known Gα interactors (i.e., Gβγ, GPCRs, effectors, RGS proteins, Ric-8A, GoLoco motifs), to specifically block GBA-dependent G protein signaling without interference with canonical signaling via GPCRs (**Fig. 5F**). Expression of GBAi reduced the efficiency of TrkB/mGluR5 crosstalk as assessed by Ca^2+^ imaging (**Fig. 5G**). This inhibitory effect was enhanced by a mutation (S252A) that increases affinity for GBA motifs but was abolished by a different mutation (W211A) that prevents GBA motif binding ^76,79^. Crucially, GBAi-S252A had no effect on the response to BDNF in cells only expressing TrkB (**Fig. S6I**), on the response to glutamate in cells co-expressing TrkB and mGluR5 (**Fig. S6J**), or on the MOR/mGluR5 crosstalk (**Fig. 5H**). Taken together with other results presented above, these data support a model in which Gβγ released upon the action of GBA proteins is responsible for the contribution of TrkB responses to the crosstalk with mGluR5 in regulating Ca^2+^ dynamics.

### G protein synergy underlies the mGluR5-dependence of BDNF-induced synaptic plasticity

To test if the G protein synergy model of TrkB/mGluR5 crosstalk is consistent with BDNF-LTP, we first used two of the aforementioned perturbations to block either Gβγ or Gα_q_-GTP. To blunt the contribution of Gβγ, we produced a Cre-dependent AAV for the membrane-tethered GRK2-CT construct and injected it into CA1 with or without co-injection of a Cre construct under the CaMKII promoter to target expression to pyramidal neurons (**Fig. S7A**). After 5-6 weeks, strong expression of GRK2-CT was observed via immunohistochemistry, including clear dendritic localization (**Fig. S7A**). Consistent with our model, GRK2-CT expression blunted BDNF-LTP to a similar degree to mGluR5 blockade or knockdown (**Fig. 6A-C**), without altering basal synaptic properties (**Fig. S7B**) or DHPG responses (**Fig. S7C, D**). GRK2-CT expression in cultured hippocampal neurons (**Fig. S7E**) also impaired the BDNF-induced increase in spine density (**Fig. 6D**). To test the contribution of Gα_q_-GTP, we applied YM-254890, which produced a small potentiation of basal fEPSP amplitude (**Fig. S7F**) and fully blocked BDNF-induced LTP (**Fig. 6E-G**). YM-254890 pre-incubation also blocked BDNF-induced spine growth in cultured hippocampal neurons (**Fig. 6H**), further supporting the G protein synergy model.

**Figure 6.**
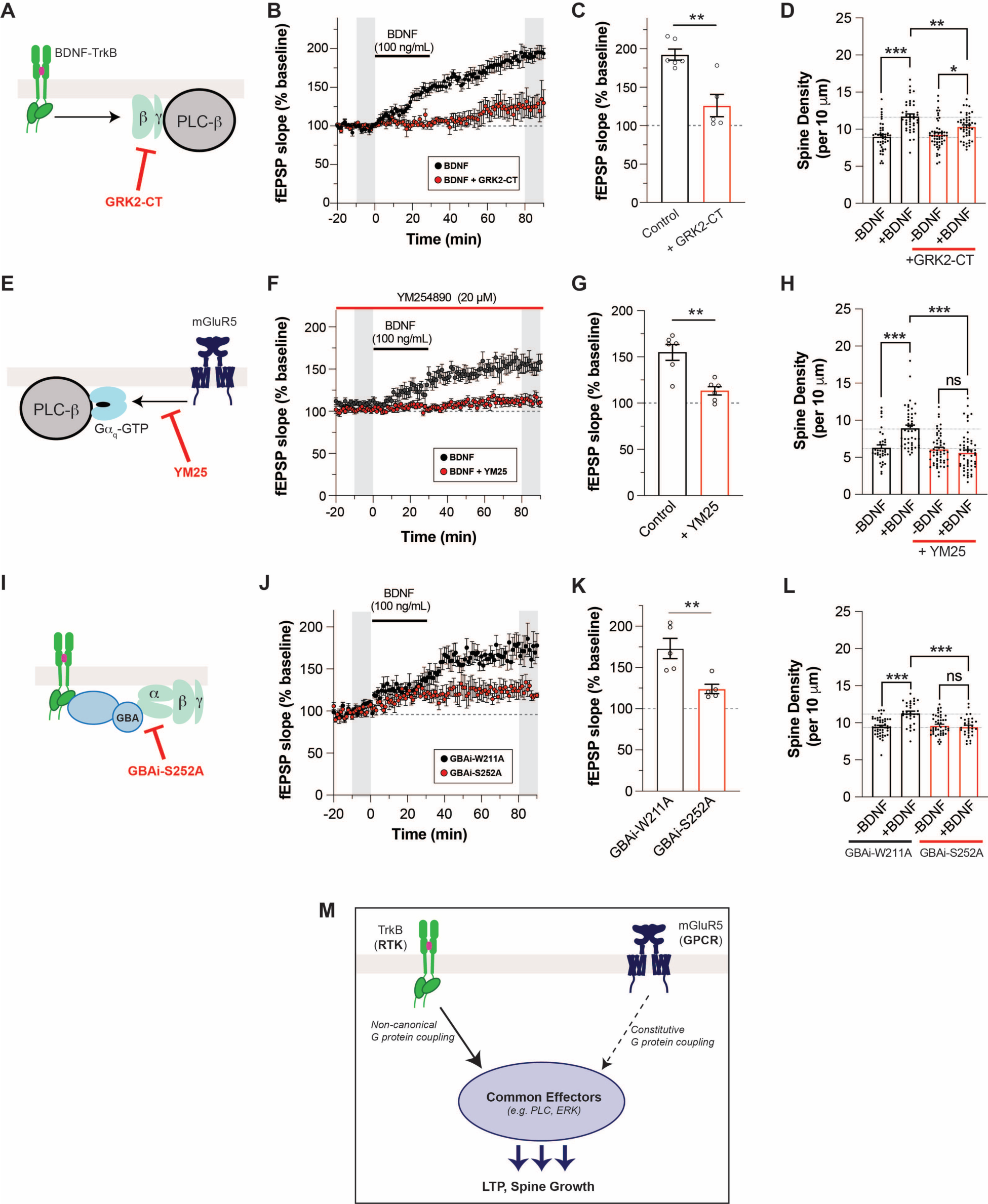
mGluR5-dependent BDNF-LTP and BDNF-induced dendritic spine growth is mediated by G protein crosstalk. **(A)** Schematic of GRK2-CT mechanism (**B-C**) fEPSP slope time course (B) showing that overexpression of GRK2-CT leads to attenuation of BDNF-LTP. In (B), grey bars show regions averaged for baseline and post-BDNF values in (C). (**D**) Bar graph summarizing the partial suppression of BDNF-induced increases in spine density with overexpression of GRK2-CT. (**E)** Schematic of YM-254890 mechanism. (**F-G**) fEPSP slope time course (F) showing that preincubation with YM-254890 leads to attenuation of BDNF-LTP. In (F), grey bars show regions averaged for baseline and post-BDNF values in (G). (**H**) Pre-incubation with YM-254890 inhibits BDNF-induced spine density increase. (**I)** Schematic of GBAi-S252A mechanism. **(J-K)** fEPSP slope time course (J) showing that overexpression of GBAi-S252A leads to attenuation of BDNF-LTP. Overexpression of the non-binding GBAi-W211A does not affect BDNF-LTP. In (J), grey bars show regions averaged for baseline and post-BDNF values in (K). (**L**) Bar graph summarizing the suppression of BDNF-induced increases in spine density with overexpression of GBAi-S252A, but not with overexpression of GBAi-W211A. (**M**) Schematic of RTK/GPCR synergy model that drives BDNF-dependent LTP and spine growth. Points in (D) and (G) represent independent slices from separate mice. Points in (D), (H), and (L) represent independent neurons from 3-4 separate preparations. Unpaired t-test is used for (C), (G), and (K). One-way ANOVA with Tukey’s multiple comparisons is used for (D), (H), and (L). All data shown as mean ± SEM; * P < 0.05, ** P < 0.01, *** P < 0.001. See also Figure S7.

Next, to test if GBA-dependent G protein signaling (**Fig. 6I**) underlies BDNF-induced plasticity, we produced Cre-dependent AAVs for GBAi-S252A or the inactive GBAi-W211A construct. Both constructs expressed well after 5-6 weeks and showed dendritic localization (**Fig. S7G**). Following transduction with GBAi-W211A AAV, normal BDNF-LTP was observed in hippocampal slices (**Fig. 6J, K**). In contrast, transduction with GBAi-S252A led to a substantial decrease in BDNF-LTP (**Fig. 6J, K**). There was no difference in basal synaptic strength between slices expressing GBAi-W211A and slices expressing GBAi-S252A (**Fig. S7H**), and no difference in DHPG-induced depression (**Fig. S7I-J)**. Lastly, expression of GBAi-S252A (**Fig. S7L**), but not GBAi-W211A (**Fig. S7K**), impaired BDNF-induced spine growth in cultured hippocampal neurons (**Fig. 6L**). Together, these results support a model in which noncanonical G protein signaling activation by GBA proteins mediates TrkB synergistic contribution to mGluR5-dependent synaptic plasticity (**Fig. 6M**).

### TrkB/mGluR5 crosstalk mechanism is generalizable across receptor subtypes

As TrkB/mGluR5 synergy merely requires simultaneous BDNF-mediated activation of TrkB and constitutive Gα_q_ activation by mGluR5, we asked if such crosstalk could proceed independently of the identity of the Gα_q_-coupled receptor. We first tested if overexpressing Gα_q_ would be sufficient to boost TrkB-induced Ca^2+^ responses. Indeed, low dose (25 ng/mL) BDNF responses were strongly enhanced in terms of both the percentage of cells responding (**Fig. 7A**) and response amplitude (**Fig. S8A**) upon Gα_q_ co-expression in HEK 293 cells. This suggests that elevated levels of Gα_q_-GTP sensitize cells to the response to TrkB activation. Consistent with this interpretation, YM25 abolished the boost in Ca^2+^ responses seen with Gα_q_ overexpression (**Fig. 7A**). We then asked if Gα_q_-coupled receptors other than mGluR5 can undergo crosstalk with TrkB. We used the TrkB-ΔPLCγ construct to abolish canonical PLC-γ mediated TrkB responses and co-expressed either mGluR1 or 5-HT_2A_R, which are both primarily coupled to Gα_q_ ^4,80^. mGluR1 co-expression allowed clear responses to BDNF in ∼20-40% of cells (**Fig. 7C, Fig. S8B**) but 5-HT_2A_R co-expression only enabled responses in ∼5-10% of cells (**Fig. 7C**). Based on our prior work ^69^, we reasoned that 5-HT_2A_R may not produce sufficient tonic Gα_q_ activation and added a subthreshold level of 5-HT (1 nM). This boosted the proportion of cells responding to BDNF to 20-30% (**Fig. 7B, C**). As an important control, the Gα_i/o_-coupled mGluR2 did not enable BDNF responses either under basal conditions or in the presence of glutamate (**Fig. 7C**), confirming the need for a Gα_q_-coupled GPCR to enable crosstalk with TrkB.

**Figure 7.**
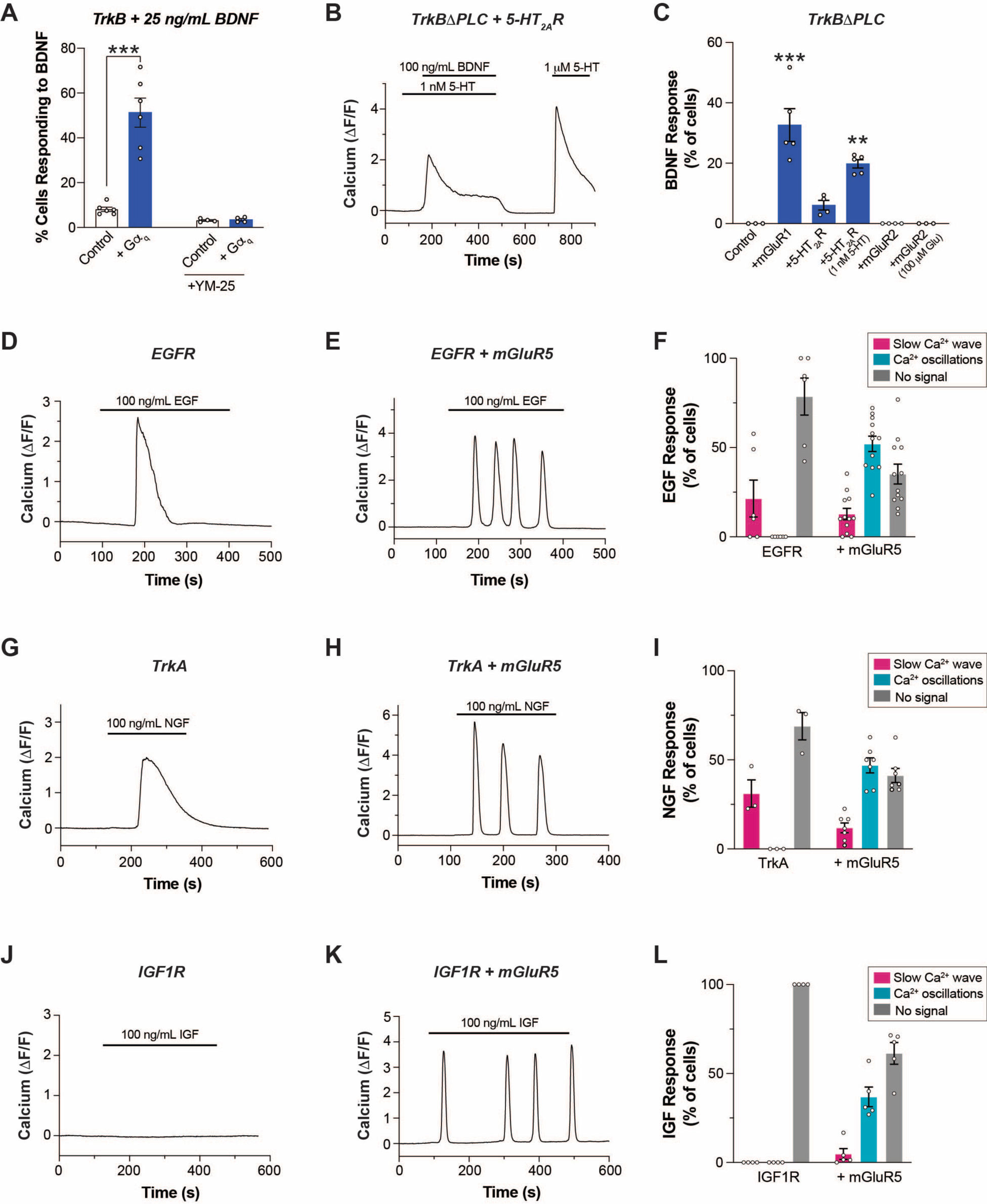
RTK/GPCR crosstalk is observed across a panel of receptors. **(A)** Summary bar graph showing that co-expression of Gα_q_ enhances Ca^2+^ responses upon low dose BDNF-induced TrkB activation in HEK 293-TrkB cells. **(B)** Representative trace showing Ca^2+^ response upon co-application of BDNF and subthreshold dose of 5-HT in HEK 293 cells co-expressing TrkB-ΔPLC and 5-HT_2A_R. **(C)** Summary bar graph showing percentage of cells responding to BDNF in HEK 293 cells co-expressing TrkBΔPLC and mGluR1 or 5-HT_2A_R or mGluR2 with or without co-application of 5-HT or glutamate. **(D-E)** Representative traces showing intracellular Ca^2+^ responses to endogenous EGFR activation by EGF in the absence (D) or presence (E) of mGluR5. **(F)** Distribution of EGF responses in the absence or presence of mGluR5 co-expression. **(G-H)** Representative traces showing intracellular Ca^2+^ responses to TrkA activation by NGF in the absence (G) or presence (H) of mGluR5. **(I)** Distribution of NGF responses in the absence or presence of mGluR5 co-expression. **(J-K)** Representative traces showing intracellular Ca^2+^ responses to IGF1R activation by IGF in the absence (J) or presence (K) of mGluR5. **(L)** Distribution of IGF responses in the absence or presence of mGluR5 co-expression. For (A) (C), (F), (I) and (L), points represent values from individual movies taken from distinct coverslips. For all conditions, data comes from at least 3 separate cell preparations. Only cells showing a response to glutamate are analyzed in the conditions with mGluR5. One-way ANOVA with Tukey’s multiple comparisons is used for (A) and (C). All data shown as mean ± SEM; *** P < 0.01, *** P < 0.001. See also Figure S8.

Given that RTKs other than TrkB have been shown to rely on heterotrimeric G proteins and/or GBA motif-mediated mechanisms to propagate signaling ^54,56,58,59,81–87^, we asked if the crosstalk observed between TrkB and Gα_q_-coupled GPCRs could be observed with other RTKs. In HEK 293 cells, we detected slow-wave Ca^2+^ responses upon EGF application, indicating signaling functionality of endogenously expressed EGFR (**Fig. 7D**). Upon mGluR5 co-expression, nearly half of the cells showed oscillatory Ca^2+^ responses to EGF (**Fig. 7E, F**), suggesting a mechanism of crosstalk analogous to that characterized for TrkB/mGluR5. Similarly, exogenous expression of TrkA, another neurotrophin receptor in the same family as TrkB ^88^, enabled nerve growth factor (NGF)-induced slow wave Ca^2+^ responses that converted to oscillatory responses upon mGluR5 co-expression (**Fig. 7G-I**). Finally, we tested the insulin growth factor 1 receptor (IGF1R) which, in contrast to other RTKs tested, showed no Ca^2+^ response to its native ligand, IGF-I (**Fig. 7J**). Strikingly, mGluR5 co-expression enabled robust IGF-I responses that were primarily oscillatory (**Fig. 7K, L)**. Co-expressions of IGF1R with mGluR1, or with 5-HT_2A_R, both also enabled IGF responses in cells expressing IGF1R (**Fig. S8C, D**). This further supports the finding that Gα_q_ tone can re-shape the acute response to RTK activation. Finally, we tested if EGF- and IGF-induced, Gα_q_-dependent Ca^2+^ responses were mediated by the same G protein synergy mechanism as observed with TrkB. Indeed, both EGF- and IGF-induced Ca^2+^ oscillatory responses upon co-expression of mGluR5 were decreased by co-expression of DN-Gα_i3_, GRK2-CT, or GBAi-S252A (**Fig. S8E, F**), suggesting a similar mechanism to TrkB/mGluR5 crosstalk. Together, these data reveal that RTK activation can synergize with tonic GPCR activation to re-shape downstream signaling dynamics.

## Discussion

Our work identifies a novel mode of RTK/GPCR crosstalk by which mGluR5 acts as a critical mediator of TrkB effects by amplifying and altering the spatiotemporal dynamics of downstream signaling to drive a unique form of BDNF-induced synaptic plasticity. We show that TrkB enhances constitutive mGluR5 activity to initiate a mode-switch leading to sustained, oscillatory Ca^2+^ signaling, as well as enhanced MAP kinase pathway activation. This crosstalk is mediated by cooperativity between Gα_q_-GTP, released by mGluR5, and Gβγ, released by non-canonical TrkB signaling. This molecular mechanism appears to be conserved across different RTKs and Gα_q_-coupled GPCRs. Our findings thus support a new “G protein synergy” model (**Fig. 5A**; **Fig. 6M**), in which simultaneous activation of both an RTK and a GPCR drives non-linear signal summation to re-shape the long-term consequences of receptor activation.

Given their extensive co-expression throughout the nervous system and the key roles of both TrkB and mGluR5 in synaptic plasticity, this neurotrophin/neurotransmitter receptor signaling crosstalk may contribute to a wide range of phenomena that have been previously attributed to either individual receptor system. For example, modulating TrkB or mGluR5 signaling has been proposed for treatment of many neurodegenerative and neuropsychiatric diseases. Recently, both TrkB and mGluR5 signaling have been shown to mediate the synaptic adaptations associated with rapid antidepressant action ^6,89,90^, which may employ a similar form of synaptic potentiation to that observed in this study. In this context, our findings that TrkB-driven synaptic plasticity is significantly enhanced by mGluR5 positive allosteric modulators (**Fig. 1E, 2D**) has potential therapeutic implications for next generation antidepressant treatment strategies that would target this form of neuromodulatory crosstalk.

Furthermore, the “G protein synergy” model proposed here represents a novel paradigm of RTK/GPCR crosstalk with relevance beyond the TrkB/mGluR5 combination and outside of the brain. We show that such crosstalk is observed with EGFR, TrkA, and IGF1R, critical RTKs with major relevance in cancer, development, and metabolism. For example, many cancers are associated with constitutively active Gα_q_ mutations ^91–93^, which may provide a means of amplifying the pro-cancer signaling of RTKs ^94,95^ and serve as promising therapeutic targets ^96^. In contrast to prior work that uncovered modes of GPCR transactivation of RTKs. ^31,32,37,38,97–101^, our study represents, to our knowledge, the first example of an RTK engaging constitutive GPCR signaling to mediate its effects by expanding its signaling repertoire. In our model, a downstream effector (i.e. PLC) serves as a coincidence detector to amplify the response to simultaneous RTK and GPCR activation. Notably, in line with this model, a recent study found that RTK-initiated downstream signaling can be blunted by Gα_q_ inhibition ^102^. Importantly, this crosstalk mode does not require direct heteromerization between RTK and GPCR, although the relative localization of the two receptors is likely a critical determinant of when and where such crosstalk may occur in native systems. In principle, this form of crosstalk could apply to many RTK/Gα_q_-coupled GPCR pairs and is likely subject to further layers of tuning and regulation, motivating future studies across biological contexts.

### Limitations of the study

This study demonstrates that BDNF-induced modes of hippocampal synaptic plasticity are dependent on coincident activity of mGluR5, a Gα_q_-coupled receptor. While we present a multitude of evidence for a G protein synergy model (**Fig. 5A**), it is important to note that many of our perturbations, such as expression of GRK2-CT, may affect G protein tone broadly and thus, perturb cells and synapses in ways that are difficult to interpret. Furthermore, other modes of crosstalk likely occur simultaneously to contribute to the complex signaling response to BDNF. For example, given that both TrkB and mGluR5 stimulate release of intracellular Ca^2+^ stores, Ca^2+^-induced Ca^2+^ release ^103^ may contribute to crosstalk in the dendritic spines and shafts where we observe complex Ca^2+^ signaling dynamics. There may also be more indirect downstream modes of RTK/Gα_q_ signaling pathway crosstalk at play, including feedback excitation and inhibition. This may explain why MPEP application blocks both BDNF-induced Ca^2+^ oscillations and slow wave Ca^2+^ responses (**Fig. 4; Fig. S5**). Furthermore, it is worth noting that both TrkB and mGluR5 undergo complex modes of endocytosis, trafficking and endosomal signaling ^104,105^, contributing to the functional interactions between receptors. Overall, the relative contribution of different modes of crosstalk likely shapes the degree and spatiotemporal dynamics of crosstalk between TrkB and mGluR5 in different contexts. This is also likely the case for tuning the crosstalk between other RTKs and other Gα_q_-coupled receptors that we propose as a general mode of receptor-receptor interaction (**Fig. 7**).

In addition to the possibility of myriad modes of signaling and trafficking crosstalk, our study also raises questions regarding RTK/G protein coupling. While our second messenger signaling and co-IP experiments point to functional coupling of TrkB with G proteins, we were unable to reliably detect G protein activation (e.g., Gβγ release) upon TrkB stimulation using bioluminescence resonance energy transfer assays established for GPCRs. This suggests that RTK-mediated G protein activation is substantially weaker than activation via bona fide GPCRs and dependent on amplification via crosstalk as seen with Ca^2+^ and cAMP imaging assays. Consistent with this idea, it has been shown that the GBA proteins presumably involved in mediating G protein signaling by RTKs, are weaker activators than GPCRs both in vitro and in cell-based assays ^57,76,79^. Ultimately, more work is needed to better understand indirect and direct modes of RTK/G protein coupling across receptor subtypes.

## Supplemental figure titles

Figure S1. Further electrophysiological analysis of BDNF-LTP in hippocampal slices, related to Figure 1.

Figure S2. Further spine imaging and dendritic calcium imaging analysis, related to Figure 1 and Figure 2.

Figure S3. Characterization of BDNF-induced TrkB calcium signaling in HEK 293 cells, related to Figure 3.

Figure S4. Further characterization of mGluR5 modulation of BDNF-induced TrkB signaling, related to Figure 3.

Figure S5. Sensitivity of BDNF-induced calcium oscillations to mGluR5 and TrkB perturbations, related to Figure 4.

Figure S6. Further analysis of G protein synergy crosstalk model, related to Figure 5.

Figure S7. Further analysis of G protein-perturbations on BDNF-driven synaptic plasticity, related to Figure 6.

Figure S8. Further analysis of G protein-perturbations on RTK/GPCR crosstalk, related to Figure 7.

## Methods

### Experimental Model and Subject Details

#### Animals

Animal care was in accordance with Weill Cornell Institutional Animal Care and Use Committee. Mice were maintained with a 12-hour light/dark cycle at 18°C - 22°C and had ad libitum access to water and food. 8–10 week-old wild-type C57BL/6J male mice (Jackson laboratory) were used for electrophysiology. The mGluR5^FL/FL^ mouse was purchased from Jackson Laboratory (B6.129-*Grm5tm1.1Jixu*/J, JAX stock #028626), and genotyping was performed per Jackson Laboratory’s protocol. All litters were weaned and sex segregated at P21. For primary hippocampal neuronal cultures, pregnant wild-type C57BL/6N mice were purchased from Charles River.

#### Human cell lines

HEK 293 cells were purchased from ATCC (CRL-1573) and authenticated by Bio-Synthesis, Inc. HEK 293-TrkB stable cell line is as previously described ^48^. All cells were tested for mycoplasma every 8 weeks with ATCC Universal Mycoplasma Detection Kit to ensure no contamination. HEK 293 cells were maintained in HEK media, consisting of DMEM (Gibco) supplemented with 10% fetal bovine serum, 100 mM sodium pyruvate, and 100 U/mL penicillin-streptomycin. HEK 293-TrkB stable cells were maintained in HEK media with 200 mg/ml G418 selection antibiotic to sustain a pure population of HEK 293-TrkB cells. Cells were passaged every 3-4 days when they reached to 90% confluency.

### Methods details

#### Reagents and antibodies

Recombinant human BDNF (Cat # 450-02), NT-3 (Cat # 450-03), NT-4 (Cat # 450-04), EGF (Cat #AF-100-15), IGF-I (Cat #100-11), NGF (Cat #450-01) were purchased from Peprotech and reconstituted in Invitrogen Ultra-Pure DNase/RNase-Free water. Picrotoxin (Cat #1128), tetrodotoxin (TTX, Cat # 1069), DHPG (Cat # 0805), MPEP (Cat # 1212), U73122 (Cat # 1268), U73343 (Cat # 4133), PD98059 (Cat # 1213), DAMGO (Cat # 1171) and baclofen (Cat # 0417), were purchased from Tocris Bioscience. YM254890 was purchased from Cayman Chemical. VU-29 was purchased from HelloBio (Cat # HB0642). Antibodies for western blots and co-immunoprecipitation assays were anti-phospho-p44/42 MAPK (Cell Signaling, Technology, 9101), anti-p44/42 MAPK (Cell Signaling Technology, 9102), anti-β-actin (Sigma, A1978), anti-TrkB (Millipore Sigma, 07-225), and anti-Gα_i3_ (Santa Cruz, H-7). Antibodies for immunocytochemistry were TrkB (R&D, AF1494), mGluR5 (Alomone, AGC-007), Alexa Fluor 546 Phalloidin (Invitrogen, A22283). Antibody for immunohistochemistry was anti-HA-Tag (Cell Signaling, C29F4, 3724) and anti-myc-Tag (Cell Signaling, 71D10, 2278).

#### Electrophysiology

Coronal brain slices (300 µm thick) containing hippocampus were prepared from male mice (C57BL/6J wildtype or mGluR5^FL/FL^). Following isoflurane anesthesia, mice were transcardially perfused, followed by brain extraction. Slices were cut with a vibratome (VT1200S, Leica) in ice-cold, oxygenated NMDG-HEPES aCSF ^106^ containing (in mM): 92 NMDG, 2.5 KCl, 1.25 NaH_2_PO_4_, 30 NaHCO_3_, 20 HEPES, 25 glucose, 2 thiourea, 5 Na-ascorbate, 3 Na-pyruvate, 0.5 CaCl_2_·2H_2_O, and 10 MgSO_4_·7H_2_O, pH to 7.3–7.4. Slices were incubated for 15-20 min on NMDG-HEPEs aCSF at 37°C then transferred to an incubation chamber at room temperature for at least 1.5 hrs containing HEPES aCSF (in mM): 92 NaCl, 2.5 KCl, 1.25 NaH_2_PO_4_, 30 NaHCO_3_, 20 HEPES, 25 glucose, 2 thiourea, 5 Na-ascorbate, 3 Na-pyruvate, 2 CaCl_2_·2H_2_O, and 2 MgSO_4_·7H_2_O, pH to 7.3–7.4. Individual slices were transferred to an immersion recording chamber, where they were submerged in oxygenated aCSF containing (in mM): 124 NaCl, 2.5 KCl, 1.25 NaH_2_PO_4_, 24 NaHCO_3_, 12.5 glucose, 5 HEPES, 2 CaCl_2_·2H_2_O, and 2 MgSO_4_,·7 H_2_O, and 10 µM picrotoxin, pH to 7.3-7.4, flowing 3 mL/min at room temperature. Field Excitatory Postsynaptic Potentials (fEPSP) were recorded through a carbon fiber electrode (Carbostar-1, Kation Scientific) placed in the *stratum radiatum* of the CA1 region. Evoked fEPSPs were elicited by Schaffer collateral stimulation with an extracellular bipolar tungsten electrode via a constant current stimulator (DS3, Constant Current Isolated Stimulator, Digitimer Ltd.) using monophasic currents of 200 µs duration. Baseline recording was obtained by stimulating the slice every 60 sec until the signal was stable using a current that elicited a 30–40% maximal response measured as the initial slope. fEPSPs were recorded in Axon Clampex 9.2 (Molecular Devices) using a Multiclamp 700b amplifier (Molecular Devices), a Digidata 1324 (Molecular Devices) with 10 kHz sampling and analyzed using Clampfit 9.2 (Molecular Devices). Following a stable baseline of ≥30 min, BDNF was applied for 30 min and recording proceeded for at least another 60 min. ANA-12 (10 μM), MPEP (40 µM), VU-29 (500 nM) were perfused for 20 min, while U73343 (5 µM), U73122 (5 µM), PD-98059 (50 µM), YM254890 (20 µM) for 60 min before and during BDNF perfusion. For DHPG-induced depression, stable baseline was established for at least 20 minutes before DHPG (50 μM) was perfused at 3 mL/min at room temperature. For paired-pulse analysis, paired stimuli with variable interstimulus intervals (25-500 ms) were applied to the Schaffer collaterals.

#### Stereotaxic viral injections for *ex vivo* slice electrophysiology

Mice were anesthetized with a ketamine and xylazine cocktail (0.1 mL per 10 g of body weight) and mounted on a stereotaxic frame (David Kopf Instruments). Viral injections were all done in the dorsal CA1 region of the hippocampus (2.25 mm AP, −2.00 ML, −1.25 mm DV from bregma), using a Nanoject II Auto-Nanoliter Injector (Drummond Scientific Company) equipped with a Nanofil syringe (World Precision Instruments). For *ex vivo* mGluR5 genetic knockout slices, 7-week-old mGluR5^FL/FL^ male mice were injected with 150 nL of AAV8-hSyn-mCherry-Cre (UNC Vector Core) or control AAV8-hSyn-mCherry (UNC Vector Core). Electrophysiology experiments were performed 5-6 weeks after injections to allow sufficient time for knockdown. To express HA-GRK2-CT, Cre-dependent AAV8-HA-GRK2-CT (UNC Vector Core, viral titer 6 x 10^12^) was pre-mixed with AAV5-CaMKII-mCherry-Cre (UNC Vector Core, viral titer 3.5 x 10^12^) at 2:1 volume ratio before injection into wildtype C57BL/6J 7-week-old male mice, at a final volume of 150 nL. For control, AAV5-CaMKII-mCherry (UNC Vector Core, viral titer 3.5 x 10^12^) was used. To express GBAi-W211A and GBAi-S252A, Cre-dependent AAV8-DIO-GBAi-W211A (UNC Vector Core, viral titer 1.5 x 10^12^) or AAV8-DIO-GBAi-S252A (UNC Vector Core, viral titer 1 x 10^12^) was pre-mixed with AAV5-CaMKII-mCherry-Cre (UNC Vector Core, viral titer 3.5 x 10^12^) at 2:1 volume ratio before injection into wildtype C57BL/6J 7-week-old male mice, at a final volume of 150 nL. Electrophysiology experiments were performed 5-6 weeks after. To validate viral expression, mice were perfused and coronal sections from fixed brains were prepared, stained for anti-HA (for HA-GRK2-CT) or anti-myc (for myc-GBAi-W211A and myc-GBAi-S252A), and imaged on the Zeiss LSM 880 confocal (refer to “Immunohistochemistry and confocal imaging” section).

#### Primary hippocampal neuronal cultures

Primary hippocampal neurons were prepared from E18 mice, as previously described ^107^, with some modifications. Briefly, bilateral hippocampi were collected and digested with 0.22 μm PES membrane-filtered custom digestion solution containing 5 mg/mL deoxyribonuclease I (Sigma, D4527/10KU), 1.5 mM CaCl_2_, 0.75 mM EDTA, 200 units of papain (Worthington, LS003127), and 2.5 mM L-cysteine (Sigma, C7352) for 5 min at 37°C. The cells were plated on autoclaved, nitric acid-washed, poly-L-lysine (Sigma) coated glass cover slips (Electron Microscopy Sciences) in Neuronal Plating Medium for 2-4 hours. They were maintained in modified Neurobasal/B27 Medium, which also had 1 mM sodium pyruvate (Gibco, 11360070) and 100 U/mL penicillin-streptomycin (Gibco, 15070063) in addition to the B27 and GlutaMAX-I supplements. On DIV 0, 4 μM cytosine-1-β-D-arabinofuranoside (Sigma, 251010) was added to limit glial proliferation.

#### Immunocytochemistry and confocal imaging of dendritic spines

DIV 21-24 primary hippocampal neurons were pre-treated with 1 μM TTX and the drug of interest (1 μM MPEP for 20 min, 500 nM VU-29 for 20 min, 10 μM ANA-12 for 1 hour, 5 μM U73122 for 1 hour, 50 μM PD98059 for 1 hour) before BDNF (50 ng/mL or 100 ng/mL) addition for 30 min at 37°C. The neurons were then washed twice with pre-warmed Neurobasal media and fixed with 4% paraformaldehyde/4% sucrose solution for 10 min at room temperature. Permeabilization and blocking were achieved with 0.1% Triton X-100 in 3% donkey serum and 3% BSA PBS solution for 30 min at room temperature. Primary antibodies were applied for 1 hour at room temperature or 4° overnight, per each antibody’s manufacturer protocols. Subtype-specific fluorophore-conjugated secondary antibodies at 1:500 dilutions were added for 30 min at room temperature. Coverslips were mounted with ProLong Gold antifade reagent (Invitrogen, P10144). Confocal imaging of the fixed coverslips was performed within 1 week of staining. Confocal imaging was performed at the Weill Cornell Medicine CLC Imaging Core Facility using Zeiss LSM 880 equipped with 32-element AiryScan detector for super-resolution imaging and 32-channel GaAsP array for spectral imaging. For spine imaging, AiryScan imaging was performed using Zeiss Plan-Apochromat 63x/1.4 Oil DIC M27 with NA 1.4 at with zoom 3.0x. Z-stack images were collected at 0.22 μm intervals and AiryScan deconvolution (Zeiss ZEN Black) was performed for each image. Spine density analysis was done by linearizing discrete secondary dendrites on ImageJ followed by manual counting of protruding dendritic spines performed blind to the experimental conditions. Approximately 40-70 μm of secondary dendrite was analyzed per neuron.

#### Calcium imaging in primary hippocampal neurons

For neuronal Ca^2+^ imaging experiments, pGP-AAV1-Syn-GCaMP8m-WPRE (Addgene, Cat # 162375, viral titer per well 1×10^8^ vg/mL) was added to the cultures at DIV 14-17. At DIV 18-23 hippocampal neuron coverslips were placed in a perfusion chamber at 33°C with a continual gravity-based perfusion of neuronal extracellular solution containing 1 μM TTX, 138 mM NaCl, 1.5 mM KCl, 5 mM HEPES, 2.5 mM CaCl_2_, 1.2 mM MgCl_2_, 10 mM glucose, pH 7.4. Imaging was performed on an Olympus IX83 microscope equipped with 488 nm laser illumination and a Hamamatsu ORCA-Flash4.0 V2 sCMOS camera imaging at 10 Hz with 42 ms exposure time. For the BDNF condition, baseline recording with neuronal extracellular solution perfusion was done for 5 minutes, followed by 100 ng/mL BDNF for 10 min, then washed out for 5 min. For the MPEP condition, baseline recording was done for 5 min, followed by 1 μM MPEP for 5 min, 1 μM MPEP and 100 ng/mL BDNF for 10 min, and washout for 5 min. For VU-29 condition, baseline recording was done for 5 minutes, followed by 500 nM VU-29 for 5 min, 500 nM + 50 ng/mL BDNF for 10 min, and washout for 5 min. Uniform 10 μm dendritic ROIs were manually drawn along each dendrite on ImageJ. Time series mean intensities were extracted for each ROI, and average background fluorescence subtracted. Photobleaching was corrected with a fitted exponential curve and intensities were normalized to a 1 min sliding median window as dF/F. To calculate the proportion of dendritic area with a BDNF response, ROIs were manually categorized as ROIs with or without signal based on the presence or absence of clear Ca^2+^ elevations in the presence of BDNF.

#### Plasmids and Molecular Cloning

Previously described ^108,109^ HA-SNAP-mGluR5 plasmids were used and modified via site-directed mutagenesis. For heterologous expression of TrkB, a SNAP-TrkB clone was made via Gibson assembly and contains an N-terminal signal sequence followed by an HA-tag and a SNAP-tag prior to full length rat TrkB. Point mutations (“kinase dead” K571N) and deletions (PLC-γ site: Δ806-821; FRS site: Δ510-513, Shc site: Δ806-821). GRK2-CAAX (# 166224) and cAMPr (# 99143) were purchased from Addgene. The PM-GRK2-CT plasmid ^110^ and PTX-S1 constructs were previously described ^111^. The GRK2-CT ORF was subcloned into a Cre-dependent AAV plasmid by restriction enzyme ligation and an HA-tag was added to the N-terminus. The G203A mutation was introduced into a rat Ga_i3_ clone (gift from D. Logothetis). An HA tag followed by a SNAP tag was introduced via Gibson assembly at the N-terminus of the rat MOR (gift from J. Broichhagen). The rat GABA_B_R clones were as previously described ^112^. All constructs were verified by sequencing. The pLIC-myc-GBAi (# 171753) and pLIC-myc-GBAi W211A (# 171754) plasmids were purchased from Addgene. The S252A mutation was introduced into the pLIC-myc-GBAi clone ^78^. The GBAi S252A and GBAi W211A were subcloned into a Cre-dependent AAV plasmid by restriction enzyme ligation.

#### HEK 293 cell transfection

For transfection and Ca^2+^ imaging and cAMP imaging, cells were plated on poly-L-lysine coated glass coverslips and transfected using Lipofectamine 2000 or 3000 (Thermo Fisher Scientific). For all transfections, 0.5 μg of DNA was used for each plasmid, except 0.3 μg of GCaMP6f and 0.7 μg of mGluR5-ΔECD. With experiments involving heterologous expression of mGluR5, cells were maintained in MPEP (1 μM) post-transfection to maintain cell health.

#### Calcium imaging in HEK 293 cells

GCaMP6f imaging was performed 24-48 hours after transfection at room temperature on either an inverted fluorescent Nikon Eclipse Ti2-E microscope equipped with an Andor Zyla 5.5 sCMOS camera or an Olympus IX83 microscope equipped with a Hamamatsu ORCA-Flash4.0 V2 sCMOS camera. All imaging was done with a 20x objective using either a 488 nm LED (Nikon microscope) or 488 nm laser (Olympus microscope). Movies were the acquired with 100 ms exposures at 0.5-1.0 Hz. Cells were continuously superfused with extracellular solution containing (in mM): 135 NaCl, 5.4 KCl, 10 HEPES, 2 CaCl_2_, 1 MgCl_2_, pH=7.4). Drugs were applied using a gravity driven perfusion system. Analysis was performed using ImageJ and NIS-Elements Advance Research 5.2.6 software. After selecting single-cell regions of interest, fluorescence intensities were quantified as dF/F by subtracting each ROI’s average resting baseline fluorescence from the first 2 min of recording before ligand addition (F_0_) from the fluorescence from any given time point (F_t_), divided by F_0_. Response latency was quantified as time from initial drug application to the first rise of response above baseline and response duration was defined as the amount of time required for fluorescence intensity to return to baseline levels after the initial rise (baseline-to-baseline). Response amplitude is the peak dF/F value and response frequency was defined as the reciprocal of averaged interval between peaks. Responses were identified as either “oscillatory” or “slow wave” based on response duration (>50 s for slow wave) and if multiple peaks were observed. At least 2 biological replicates (i.e. separate transfections) were included for all experiments (see Figure legends for details). All analyses were manually performed on Microsoft Excel, with statistical analysis performed on GraphPad Prism. Analysis of surface expression of SNAP-tagged constructs was performed as previously described ^113^.

#### cAMP imaging in HEK 293 cells

cAMPr imaging was performed 24-48 hours after transfection at room temperature on an inverted fluorescent Nikon Eclipse Ti2-E microscope equipped with an Andor Zyla 5.5 sCMOS camera. Briefly, movies were the acquired with 100 ms exposures at 0.25-0.5Hz with 20x objective. Cells were continuously superfused with extracellular solution same as used in calcium imaging experiments. BDNF or DAMGO was applied to cells 2 min prior to Forskolin co-application. Fluorescence intensity was analyzed in a similar way as calcium imaging. Area under the curve (AUC) of the normalized traces was used to quantify the cAMP response. Analysis was performed using ImageJ and NIS-Elements Advance Research 5.2.6 software. All analyses were manually performed on Microsoft Excel, with statistical analysis performed on GraphPad Prism

#### Western blot and Immunoprecipitation assays

For western blot lysate preparation, HEK 293-TrkB cells were treated with drugs prior to cell lysis with cold Pierce RIPA lysis buffer (Thermo Scientific, 89901) supplemented with protease inhibitor set I (Sigma, 539131) and phosphatase inhibitor set II (Sigma, 524625). Protein extracts were quantified using Pierce BCA protein assay kit (Thermo Scientific, 23225). 20 μg (HEK 293-TrkB) of protein was reduced in NuPAGE LDS sample buffer (Invitrogen, NP0007) with NuPAGE sample reducing agent (Invitrogen, NP0009) and boiled at 65°C for 10 min. Proteins were then run on 4-12% Bolt-Tris Plus mini gels for separation and transferred onto Bio-Rad Immun-Blot PVDF membrane using Bio-Rad Trans-Blot Turbo Transfer System. Membranes were blocked for at least 1 hour at room temperature prior to incubation with primary antibodies 4°C overnight or 1 hour at room temperature, depending on each antibody’s manufacturer recommendations. Following 3 washes in TBST, membranes were incubated with appropriate HRP secondary antibodies (Jackson laboratories) at room temperature for 30 minutes prior to visualization by ECL detection (Thermo Scientific, 32106) and Bio-Rad ChemiDoc XRS+ imaging system with Image Lab Software. Membranes were stripped with Restore stripping buffer (Thermo Scientific, 21063), rinsed 4 times with TBST, before blocking and probing again with other antibodies. Each experiment was repeated at least three times. Blots were analyzed on ImageJ by measuring intensity of phosphorylated ERK1/2 bands with corresponding ERK1/2 bands for normalization. For immunoprecipitation assays, cells were treated with drugs prior to cell lysis with immunoprecipitation lysis buffer (20 mM HEPES, 5 mM Mg-acetate, 125 mM K-acetate, 0.4% Triton X-100, 1mM dithiothreitol, 100 μM sodium orthovanadate, and phosphatase/protease inhibitors from Millipore Sigma as noted previously). Sepharose protein G beads were pre-incubated with antibodies to form antibody-bead complexes overnight at 4°C overnight. These complexes were then added to protein lysates for 2-4 hours at room temperature to allow for binding of desired protein to the beads. The beads were then washed 4 times in immunoprecipitation lysis buffer, then similarly reduced in NuPAGE LDS sample buffer and NuPAGE sample reducing agent prior to boiling at 65°C for 10 min to allow elution of the proteins off the beads. The immunoprecipitated proteins were then resolved as per western blot protocol above.

#### Immunohistochemistry and confocal imaging

Whole brain coronal sections (40 μm) were prepared using a sliding microtome. Serial sections were washed in TBS, incubated for 30 min in a blocking solution containing 4% normal horse serum (vol/vol), 1% BSA in TBS with 0.2% Triton X-100, and incubated overnight at 4°C with primary antibodies diluted in blocking solution. Sections were then washed in TBS and incubated for 2 hours with subtype-specific Alexa-488 conjugated secondary antibody at room temperature. After washing three times for 10 min, sections were mounted, coverslipped with water soluble glycerol-based mounting medium containing DAPI. Confocal imaging was performed on a Zeiss LSM 880 with Zeiss Plan-Apochromat 20x/0.8 DIC-II. Tiled Z-stack images were collected at 0.94 μm intervals.

#### Quantification and statistical analysis

All data are presented as means ±SEM and analyzed with GraphPad Prism 9.0 software. Statistical significances were calculated via unpaired student t test (for two group comparisons), one-way ANOVA with Šidák and Tukey’s post hoc tests to control for multiple comparisons (for three or more group comparisons), or two-way ANOVA with Sidak’s multiple comparison tests (to assess statistical significance between means), as indicated within individual figure legends. For frequency distribution histogram statistics, exact sum-of-squares F-test was used. In figures, asterisks denote statistical significance marked by * P < 0.05, ** P < 0.01, *** P < 0.001, and “n.s.” indicates no statistical significance.

## Acknowledgements

We are grateful to David Simon (Weill Cornell Medicine) for access to his Nikon Ti2-E microscope and valuable discussions. We thank Hermany Munguba (Weill Cornell Medicine) for setting up the electrophysiology rig. We thank Rasmus Herlo Jensen (Columbia University) and Logan Grosenick (Weill Cornell Medicine) for their insights on neuronal calcium signaling analysis. Lastly, we thank research technicians Roshelle Smith (Weill Cornell Medicine), Chienchun (Ted) Huang (Weill Cornell Medicine), and Rui Rong Yang (Weill Cornell Medicine) for assistance with preparation of cultures, mouse colony management, and immunohistochemistry. Funding sources: National Institute of Health grants R01NS126590 (JL, FSL), R35GM124731 (JL), R01MH123154 (FSL), R01GM136132 (MGM), Rohr Family Research Scholar Award (JL), Monique Weill-Caulier Award (JL), Pritzker Neuropsychiatric Disorders Research Consortium (FSL), NewYork–Presbyterian Center for Youth Mental Health (FSL), National Research Foundation (NRF) of Korea grant NRF-2021R1I1A3055750 (MS), Burroughs Wellcome Fund - Weill Cornell Physician-Scientist Academy Award (JK).

## Author contributions

Conceptualization: CLP, GX, JK, MGM, JL, FSL. Methodology: CLP, GX, JK, MS. Investigation: CLP, GX, JK, IS, AF, DG, PK, MS, DK. Formal Analysis: CLP, GX, JK, JL. Funding acquisition: JK, MGM, MS, JL, FSL. Project administration: JL, FSL. Supervision: JL, FSL. Writing – original draft: JL. Writing – review & editing: CLP, GX, JK, MS, MGM, JL, FSL

## Materials and data availability

All materials can be provided pending a completed material transfer agreement. Requests should be submitted to the lead contact Dr. Francis S. Lee. All data are available in the main text or the supplementary materials. This paper does not report original code.

