## Supplemental Figures for "Synaptic plasticity via receptor tyrosine kinase/G protein-coupled receptor crosstalk"

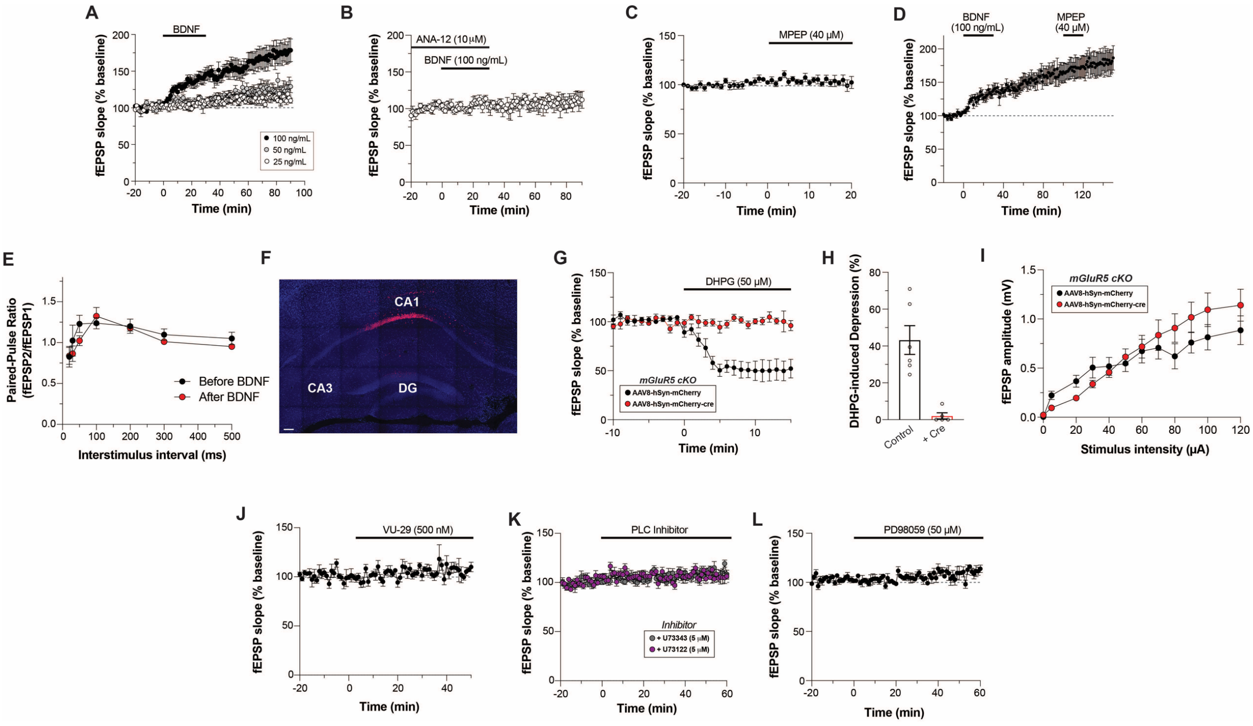


**Figure S1. Further electrophysiological analysis of BDNF-LTP in hippocampal slices. Related to Figure 1.** (**A**) fEPSP slope time course in response to various doses of BDNF for 30 min (from Time = 0 to 30 min). 100 ng/mL (n = 8 slices), 50 ng/mL (n = 6 slices), 25 ng/mL (n = 5 slices). (**B**) fEPSP slope time course with ANA-12 alone showing no effects on baseline plasticity (n = 4 slices). (**C**) fEPSP slope time course with MPEP alone showing no effects (n = 5 slices). (**D**) fEPSP slope time course showing no effect of MPEP after BDNF-LTP induction and maintenance for 60 min (n = 6 slices). (**E**) Paired-pulse ratio of fEPSPs across a range of interpulse intervals before and after BDNF 100 ng/mL perfusion for 30 min (n = 6 slices). (**F**) Representative image of dorsal CA1 hippocampal expression of AAV8-hSyn-mCherry-Cre in mGluR5$^{FL/FL}$ mice (CA1, CA3 = cornu ammonis 1, 3; DG = dentate gyrus, scale bar 100 um). (**G-H**) DHPG application decreases fEPSP slope in control mGluR5$^{FL/FL}$ slices with AAV8-hSyn-mCherry, but not when mGluR5 is knocked out with AAV8-hSyn-mCherry-Cre. Individual points in (H) denote independent slices taken from distinct mice, and unpaired student t-test is used. (**I**) Input/output curve of fEPSP amplitude under basal conditions in mGluR5$^{FL/FL}$ mice with AAV8-hSyn-mCherry (n = 4 slices) or AAV8-hSyn-mCherry-Cre (n = 5 slices). (**J-L**) fEPSP slope time courses showing no effects of VU-29 (J, n = 6 slices), PLC inhibitor U-73122 or inactive analog U-73343 alone (K, n = 6 slices for each drug), or MEK inhibitor PD-98059 alone (L, n = 6 slices). For (A-L), each slice is taken from distinct mice. All data are shown as mean ± SEM.

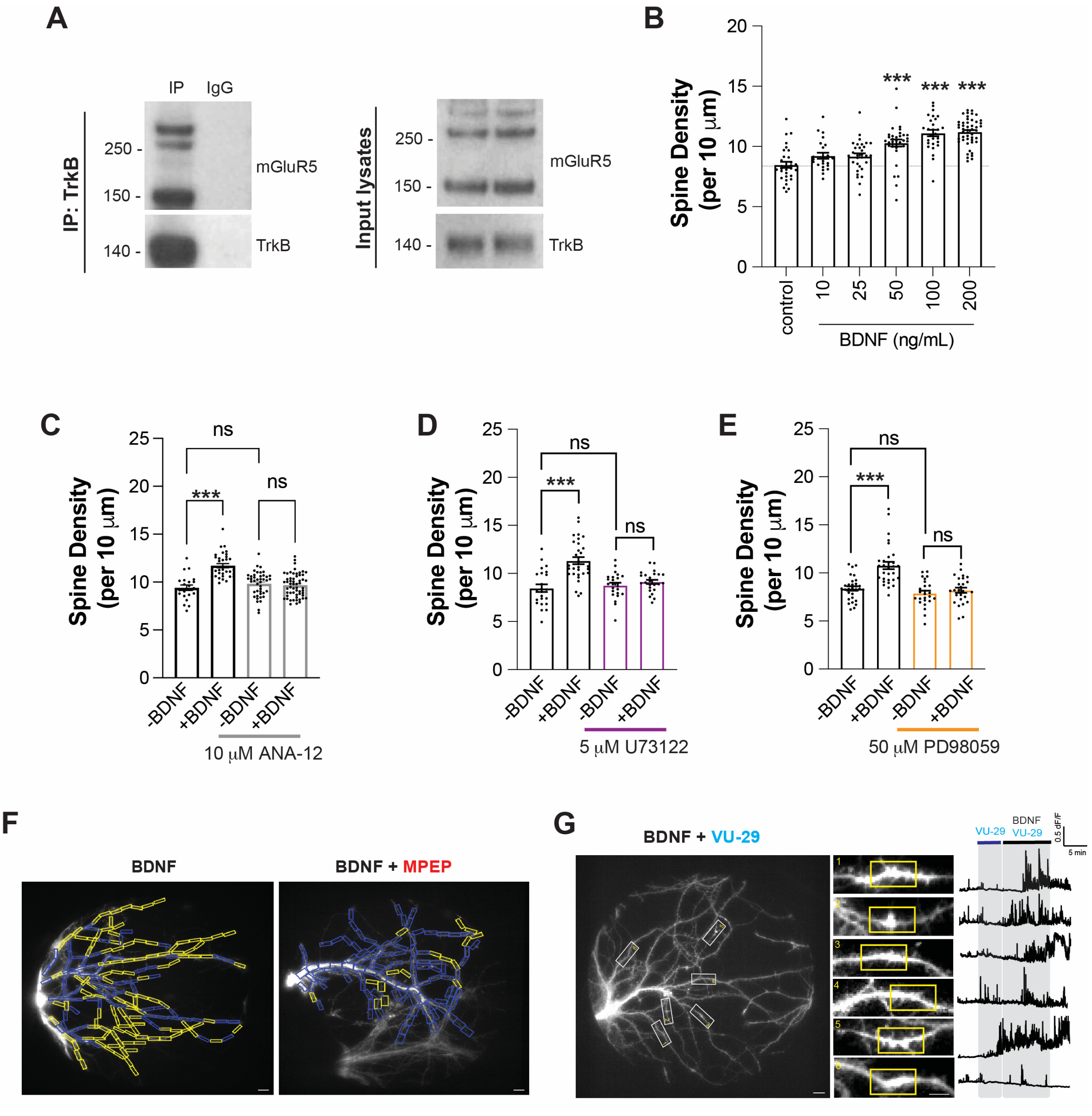


**Figure S2. Further spine imaging and dendritic calcium imaging analysis. Related to Figure 1 and Figure 2.** (**A**) Co-immunoprecipitation of endogenous TrkB and mGluR5 in DIV 19 primary hippocampal neuronal lysates. (**B**) Bar graphs showing dendritic spine density with increasing concentrations of BDNF for 30 min. (**C-E**) Bar graphs showing dendritic spine density without or with 100 ng/mL BDNF for 30 min with various pharmacological modulations. Pre-treatment with TrkB antagonist ANA-12 (C), PLC inhibitor U-73122 (D), or MEK inhibitor PD-98059 (E) for 60 min before BDNF addition prevents the BDNF-induced increase in spine density without altering basal spine density. For (B-E), individual points represent separate neurons, and data comes from two (B) or three (D-E) separate culture preparations for each graph. One-way ANOVA with Tukey's multiple comparisons is used. All data shown as mean ± SEM; ** $P < 0.01$, *** $P < 0.001$. (**F**) Representative images of hippocampal neurons showing ROIs with $Ca^{2+}$ response (yellow rectangle, 10 μm) or without $Ca^{2+}$ response (blue rectangle, 10 μm) when BDNF is perfused without and with MPEP (1 μM). (**G**) Left, representative image of hippocampal neuron expressing GCaMP8m (left, scale bar 10 μm), with snapshots of dendrites (white rectangle) during VU-29 (500 nM) and 50 ng/mL BDNF perfusion (middle, scale bar 5 μm). Right, representative trace from 10 μm ROIs with $Ca^{2+}$ response (yellow rectangle) from each dendrite.

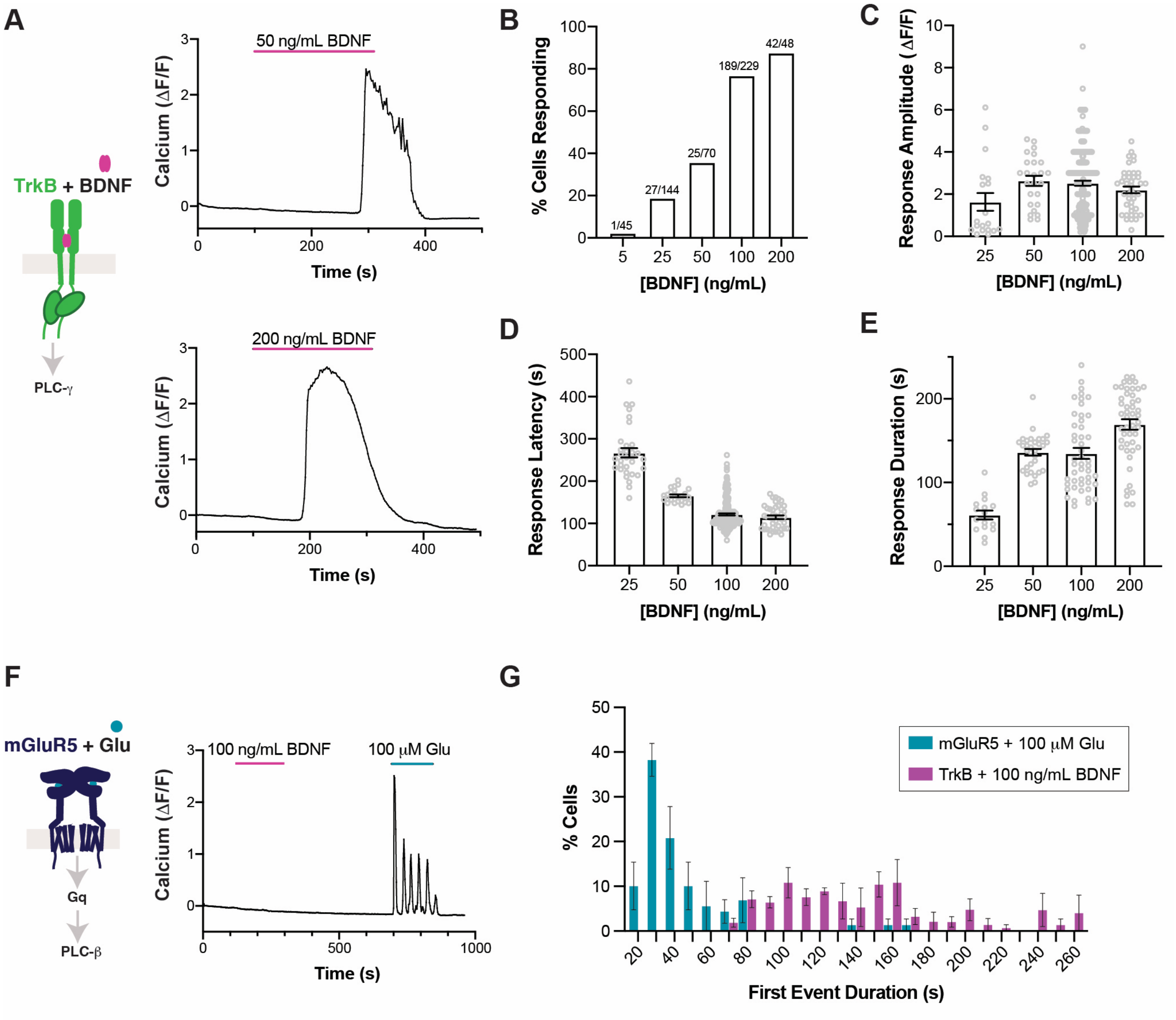


**Figure S3. Characterization of BDNF-induced TrkB calcium signaling in HEK 293 cells. Related to Figure 3.** (**A**) Representative single-cell traces showing BDNF-induced $Ca^{2+}$ responses in HEK 293-TrkB cells. (**B-E**) Bar graphs showing dose dependence of percentage of cells responding to BDNF (B), and the amplitude (C), latency (D), and duration (E) of responses. (**F**) Representative single-cell trace from a HEK 293 cell expressing mGluR5 only, showing no BDNF-induced $Ca^{2+}$ response but a clear glutamate-induced oscillatory response. (**G**) Frequency distribution histogram of the percentage of cells with duration of the first $Ca^{2+}$ event induced either by BDNF on HEK 293-TrkB cells or glutamate on HEK 293 cells transfected with mGluR5 only. The two distributions are significantly disparate by F test (34.95, Dfn 3, Dfd 20; $p < 0.0001$). For (C-E): individual points are individual cells, and data on each graph are from 3 separate cell preparations. One-way ANOVA with Tukey's multiple comparisons is used. All data shown as mean ± SEM.

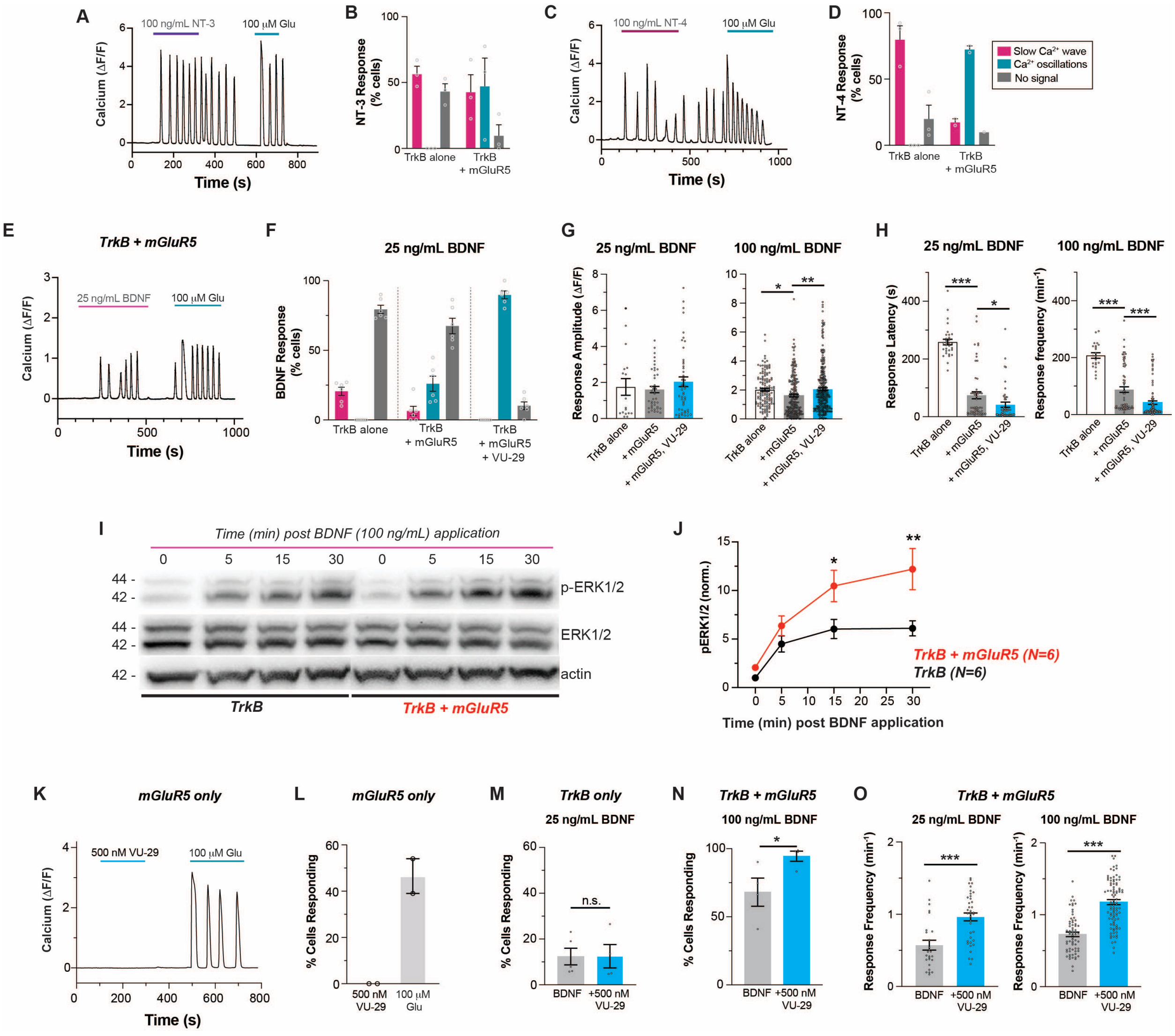


**Figure S4. Further characterization of mGluR5 modulation of BDNF-induced TrkB signaling, related to Figure 3.** (**A-D**) Representative traces and summary bar graphs showing NT-3-induced oscillatory $Ca^{2+}$ responses (A-B) and NT-4 induced oscillatory $Ca^{2+}$ responses (C-D) in HEK 293-TrkB cells expressing mGluR5. (**E**) Representative trace showing $Ca^{2+}$ oscillations induced by low dose BDNF in HEK 293-TrkB cells transfected with mGluR5. (**F**) Distribution of BDNF-induced $Ca^{2+}$ response with low dose of BDNF in HEK 293-TrkB cells with and without mGluR5 co-expression, as well as with and without co-application of 500 nM VU-29. (**G-H**) Bar graphs showing the effects of mGluR5 and 500 nM VU-29 on low dose (25 ng/mL) and high dose (100 ng/mL) BDNF response amplitude (G) and latency (H). (**I-J**) Representative western blot (I) and BDNF application time course graph (J) showing phospho-ERK1/2 response to BDNF in HEK 293-TrkB cells without and with mGluR5 co-expression. (n = 6 individual blots). (**K-L**) Representative trace (K) and summary bar graph (L) showing that 500 nM VU-29 alone does not elicit $Ca^{2+}$ responses in HEK 293 cells expressing mGluR5 only. (**M**) Bar graph showing lack of any effect of co-application of VU-29 on percentage of cells responding to low dose BDNF (25 ng/mL) in HEK 293-TrkB cells without mGluR5 co-expression. (**N**) Bar graph showing effects of co-application of VU-29 with BDNF in HEK 293-TrkB cells with mGluR5 co-expression. Only cells responding to glutamate were included in the graph. (**O**) Bar graphs showing that VU-29 increases the frequency of low dose and high dose BDNF-induced $Ca^{2+}$ oscillations in HEK293-TrkB cells with co-expression of mGluR5. For (B), (D), (F), (L), (M), and (N): individual points come from separate coverslips. For (G), (H), and (O): individual points come from separate cells. For all conditions, data comes from at least 3 separate cell preparations. Two-way ANOVA with Sidak's multiple comparison test is used for (J). One-way ANOVA with Tukey's multiple comparison is used for (G) and (H). Unpaired t-test is used for (M), (N) and (O). All data shown as mean ± SEM; * $P < 0.05$, ** $P < 0.01$, *** $P < 0.001$.

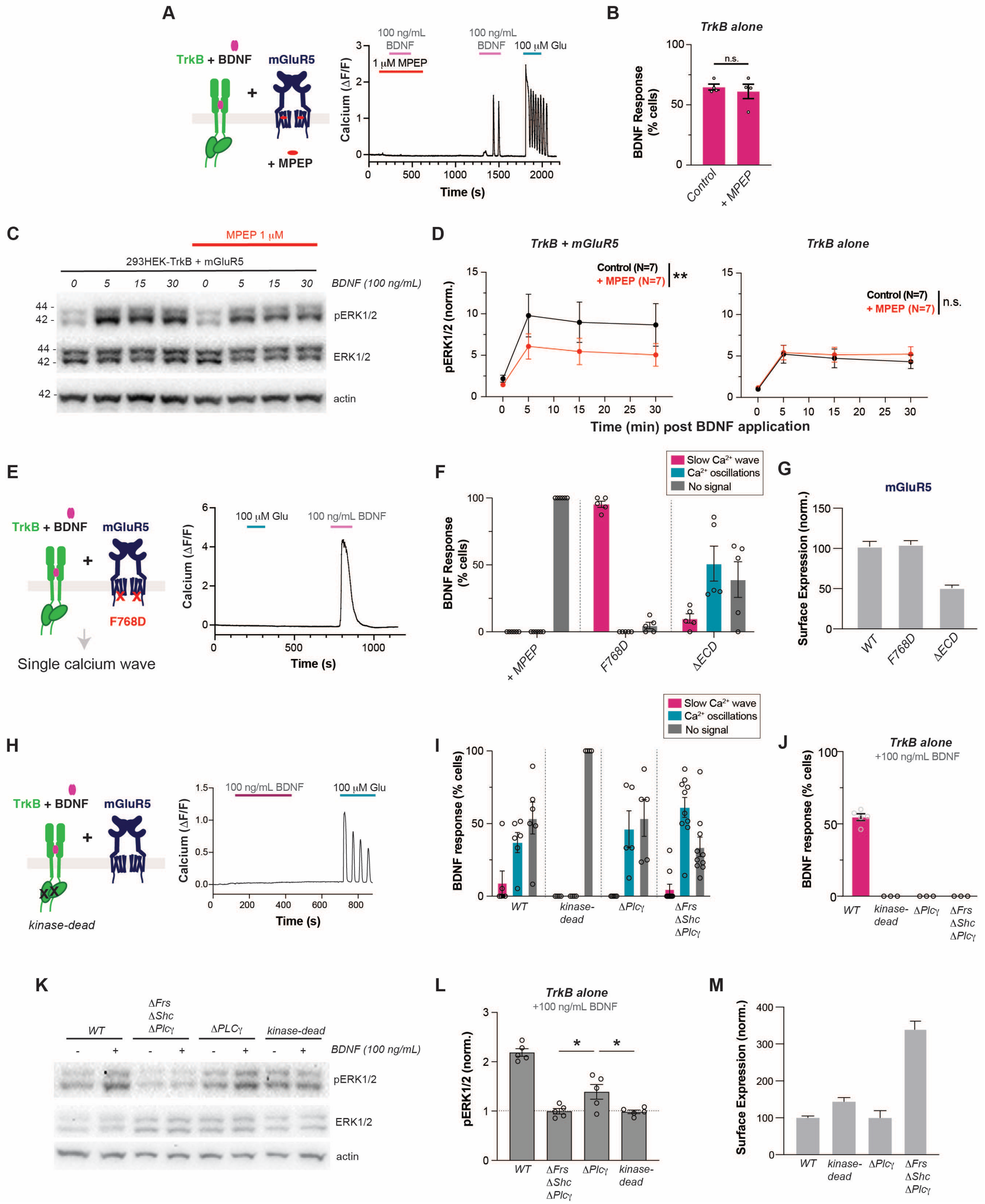


**Figure S5. Sensitivity of BDNF-induced calcium oscillations to mGluR5 and TrkB perturbations, related to Figure 4.** (**A**) Representative $Ca^{2+}$ trace showing MPEP blocking BDNF-induced $Ca^{2+}$ oscillations, followed by induction of BDNF-induced and glutamate-induced $Ca^{2+}$ oscillations after MPEP washout. (**B**) Bar graph showing that co-application of MPEP does not change the percentage of cells responding to BDNF in HEK293-TrkB cells without mGluR5 co-expression. (**C-D**) Representative western blot (C) and quantification (D) of BDNF-induced ERK activation in HEK 293-TrkB cells co-expressing mGluR5, showing a reduced response in the presence of MPEP. (**E**) Representative $Ca^{2+}$ signaling trace with lack of BDNF and glutamate-induced $Ca^{2+}$ oscillations in HEK 293 cells expressing mGluR5 F768D, a mutant mGluR5 with reduced affinity for G protein coupling. (**F**) Distribution of BDNF-induced $Ca^{2+}$ responses in HEK 293-TrkB cells expressing mGluR5 WT with 1 μM MPEP, mGluR5 F768D, and mGluR5ΔECD. (**G**) Bar graph showing normalized surface expression for mGluR5 WT, mGluR5 F768D, and mGluR5ΔECD. (**H**) Representative trace showing the lack of BDNF-induced $Ca^{2+}$ response in HEK 293 cells expressing TrkB kinase-dead and mGluR5, with maintained mGluR5 response to glutamate. (**I**) Distribution of $Ca^{2+}$ response types in HEK 293 cells co-expressing mGluR5 with wild type or mutant TrkB. (**J**) Bar graph showing percentage of cells responding to BDNF in HEK 293 cells transfected with wild type or mutant TrkB. (**K**-**L**) Representative western blot (**K**) and quantification (**L**) of BDNF-induced ERK activation in HEK 293 cells transfected with TrkB wild type and mutants. BDNF (100 ng/mL) stimulation was done for 15 min. Quantification of pERK/ERK ratio shows BDNF-induced ERK activation as compared to each TrkB mutant baseline condition. (**M**) Bar graph showing the surface expression of TrkB mutants normalized to surface expression of wild type TrkB. For (B), (F), (I) and (J): individual points come from separate coverslips. For (L): individual points represent separate western blots with lysates prepared from disparate cell preparations. For all conditions, data comes from at least 3 separate cell preparations. Unpaired t-test is used for (B). Two-way ANOVA is used for (D). One-way ANOVA with Sidak's multiple comparisons is used for (L). All data shown as mean ± SEM; * $P < 0.05$, ** $P < 0.01$.

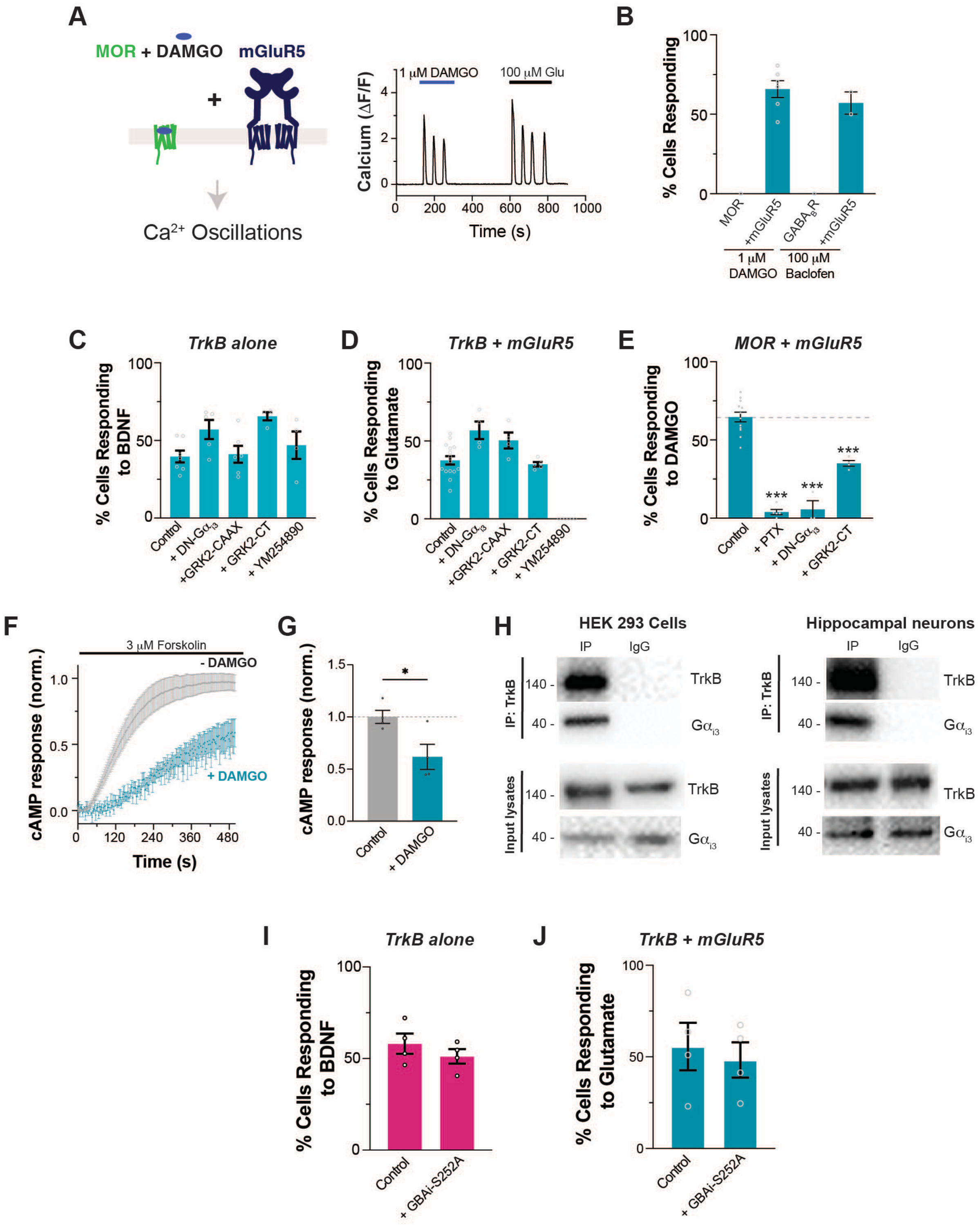


**Figure S6. Further analysis of G protein synergy crosstalk model, related to Figure 5.** (**A**) Representative trace showing DAMGO-induced $Ca^{2+}$ oscillations in HEK 293 cells co-expressing MOR and mGluR5. (**B**) Summary bar graph of percentage of cells with agonist- induced $Ca^{2+}$ response in HEK 293 cells expressing MOR or $GABA_BR$ without or with mGluR5. (**C**) Summary bar graph showing the lack of effects of G protein perturbations and 20 μM YM254890 on $Ca^{2+}$

response to BDNF in HEK 293-TrkB cells without mGluR5 co-expression. (**D**) Summary bar graph showing the percentage of HEK 293-TrkB cells with mGluR5 co-expression showing glutamate-induced $Ca^{2+}$ responses. G protein perturbations show no clear effect on glutamate response, except for YM254890 (20 μM). (**E**) Summary bar graph showing decreased responses to DAMGO in HEK 293 cells co-expressing MOR and PTX, DN-$G\alpha_{i3}$, or GRK2-CT with mGluR5. (**F-G**) Normalized traces (F) and summary bar graph (G) showing forskolin-induced cAMP response in HEK 293 cells expressing MOR with or without co-application of 1 μM DAMGO. (**H**) Representative co-immunoprecipitation blots of TrkB and $G\alpha_{i3}$ protein in HEK293-TrkB cells and primary hippocampal neurons. (**I-J**) Summary bar graph showing the effects of co-expression of GBAi-S252A on percentage of cells responding to BDNF in HEK293-TrkB cells without mGluR5 co-expression (I) or glutamate in HEK293-TrkB cells with mGluR5 co-expression (J). For (B), (C), (D) (E), (G), (I) and (J): Individual points come from separate coverslips from at least 3 separate cell preparations. One-way ANOVA with Dunnett's multiple comparison test is used to compare with control group for (C), (D), and (E). Unpaired t-test was used for (G), (I), and (J). All data shown as mean ± SEM; * $P < 0.05$, *** $P < 0.001$.

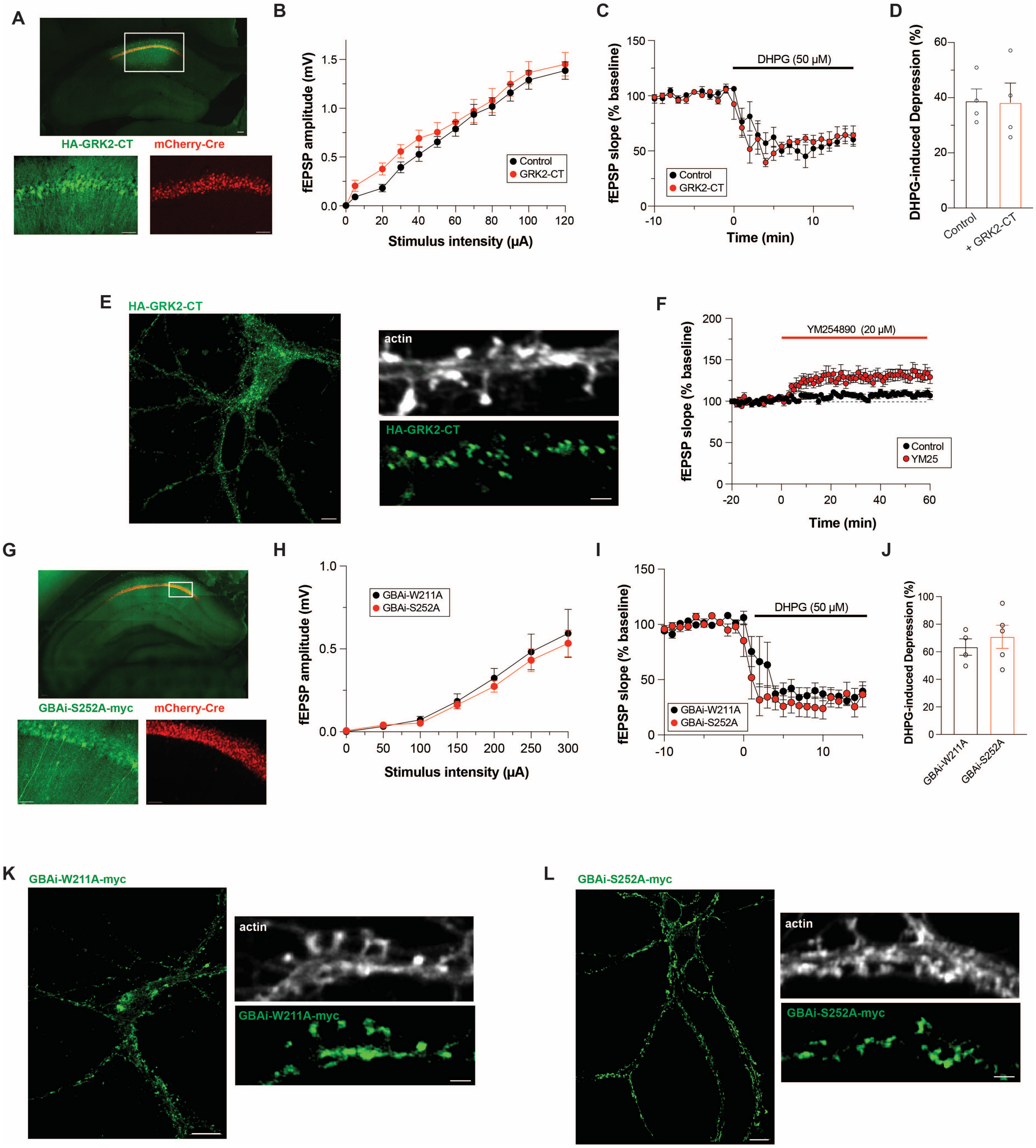


**Figure S7. Further analysis of G protein-perturbations on BDNF-driven synaptic plasticity, related to Figure 6.** (**A**) Representative confocal image of the hippocampus, showing expression of CaMKII-mCherry-Cre and HA-GRK2-CT in the dorsal CA1 (top, scale bar 100 μm). Close-up of the dorsal CA1 region (top, white rectangle) shown with soma and dendritic filling of the HA-GRK2-CT and mCherry-Cre (bottom, scale bar 50 μm). (**B**) Input/output curve of fEPSP amplitude showing no change in basal synaptic strength upon expression of HA-GRK2-CT (n = 4 slices). (**C-D**) fEPSP slope time course (C) and summary bar graph (D) showing no effects of HA-GRK2-

CT expression on DHPG induced synaptic inhibition. Individual points represent independent slices taken from distinct mice. (**E**) Representative confocal image of HA-GRK2-CT expression in primary hippocampal neurons (left, scale bar 5 μm) with a zoom-in of the dendritic shaft (right, scale bar 1 μm). (**F**) Incubation with YM-254890 elicits a modest increase in fEPSP slope that stabilizes within 20 min (n = 6 slices from distinct mice). (**G**) Representative confocal image of the hippocampus, showing expression of GBAi-S252A-myc and CaMKII-mCherry-Cre in the dorsal CA1 (top, scale bar 100 μm). Close-up of the dorsal CA1 region (top, white rectangle) shown with soma and dendritic filling of the GBAi-S252A-myc (bottom, scale bar 50 μm). (**H**) Input/output curve of fEPSP amplitude in slices injected with GBAi-W211A (n = 6 slices) and GBAi-S252A (n = 5 slices). (**I-J**) fEPSP slope time course (I) showing that DHPG application decreases fEPSP slope in slices expressing GBAi-W211A (n = 4 slices) and GBAi-S252A (n = 5 slices). In (I), grey bars show regions averaged for baseline and post-BDNF values in (J). For (D) and (J), unpaired t-test is used. All data shown as mean ± SEM.

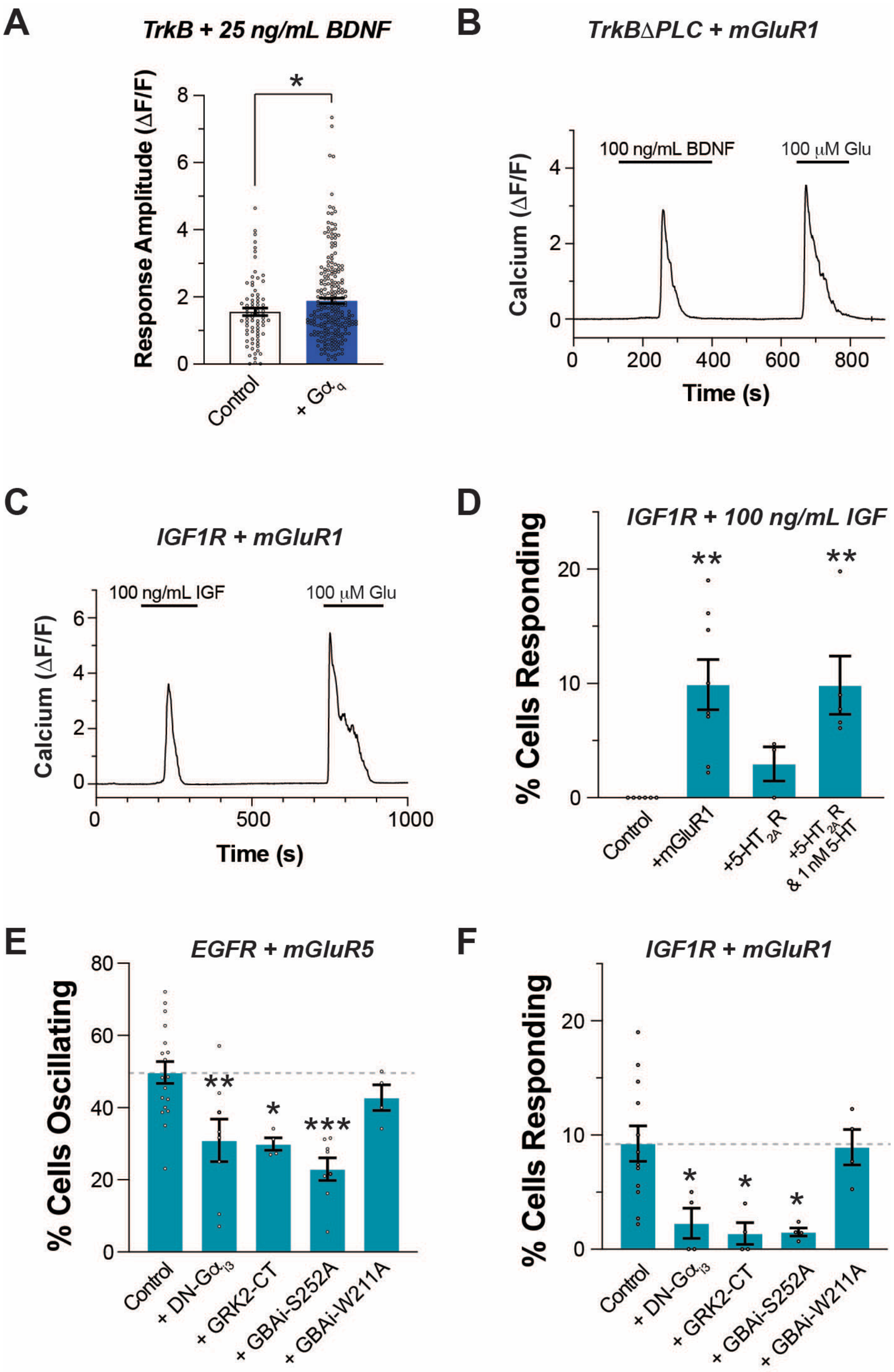


**Figure S8. Further analysis of G protein-perturbations on RTK/GPCR crosstalk, related to Figure 7.** (**A**) Bar graph showing the effects of $G\alpha_q$ co-expression on low dose BDNF-induced calcium response amplitude in HEK293-TrkB cells. (**B**) Representative trace showing BDNF-induced $Ca^{2+}$ response in HEK 293 cells co-expressing TrkB-ΔPLCγ (B) or $IGF_1R$ (C) and mGluR1. (**D**) Summary bar graph showing percentage of cells responding to IGF in HEK 293 cells co-expression of $IGF_1R$ and mGluR1 or $5\text{-}HT_{2A}R$. (**E**) Summary bar graph showing decreased responses to EGF in HEK 293 cells co-expressing mGluR5 and DN-$G\alpha_{i3}$, GRK2-CTor GBAi-

S252A but not GBAi-W211A. (**F**) Summary bar graph showing decreased responses to IGF in HEK 293 cells co-expressing $IGF_1R$, mGluR1 and DN-$G\alpha_{i3}$, GRK2-CT, or GBAi-S252A but not GBAi-W211A. Only cells responding to glutamate or 5-HT were included for (D), (E) and (F). Unpaired t-test is used for (A). One-way ANOVA with Dunnett's multiple comparison test is used to compare with control group for (D), (E) and (F). All data shown as mean ± SEM; * $P < 0.05$, *** $P < 0.001$.
